# Pumilio Proteins Exert Distinct Biological Functions and Multiple Modes of Post-Transcriptional Regulation in Embryonic Stem Cell Pluripotency and Early Embryogenesis

**DOI:** 10.1101/751909

**Authors:** Katherine E. Uyhazi, Yiying Yang, Na Liu, Hongying Qi, Xiao A. Huang, Winifred Mak, Scott D. Weatherbee, Xiaoling Song, Haifan Lin

## Abstract

Gene regulation in embryonic stem cells (ESCs) has been extensively studied at the epigenetic-transcriptional levels, but not at the post-transcriptional levels. Pumilio (Pum) proteins are among the few known translational regulators required for stem cell maintenance in invertebrates and plants. Here we report the essential function of two murine Pum proteins, Pum1 and Pum2, in ESCs and early embryogenesis. Pum1/2 double mutants are developmentally delayed at the morula stage and lethal by embryonic day 8.5 (e8.5). Correspondingly, Pum1/2 double mutant ESCs display severely reduced self-renewal and differentiation, revealing the combined function of Pum1 and Pum2 in ESC pluripotency. Remarkably, Pum1-deficient ESCs show increased expression of pluripotency genes but not differentiation genes, indicating that Pum1 mainly promote differentiation; whereas Pum2-deficient ESCs show decreased expression of pluripotency genes and accelerated differentiation, indicating that Pum2 promotes self-renewal. Thus, Pum1 and Pum2 each uniquely contributes to one of the two complementary aspects of pluripotency. Furthermore, we show that Pum1 and Pum2 achieve ESC functions by forming a negative auto- and inter-regulatory feedback loop that directly regulates at least 1,486 mRNAs. Pum1 and Pum2 regulate target mRNAs not only by repressing translation as expected but also by promoting translation and enhancing or reducing mRNA stability of different target mRNAs. Together, these findings reveal the distinct roles of individual mammalian Pum proteins in ESCs and their collectively essential functions in ESC pluripotency and embryogenesis. Moreover, they demonstrate three novel modes of regulation of Pum proteins towards target mRNAs.

**SIGNIFICANCE STATEMENT:** This report demonstrates the essential functions of mammalian Pumilio (Pum) proteins for embryonic stem cells (ESCs) pluripotency and embryogenesis. Moreover, it reveals the contrasting but complementary function of individual Pum proteins in regulating distinct aspects of ESC pluripotency, despite their largely overlapping expression and extremely high homology. Furthermore, it unravels a complex regulatory network in which Pum1 and Pum2 form a negative auto- and inter-regulatory feedback loop that regulates 1,486 mRNAs not only by translational repression as expected but also by promoting translation and enhancing or reducing stability of different target mRNAs, which reveals novel modes of post-transcriptional regulation mediated by Pum.

## INTRODUCTION

Embryonic stem cells (ESCs) possess dual abilities to self-renew and to differentiate into any cell type in the body. Recent work has identified transcription factors that are essential for stem cell self-renewal and pluripotency, but the role of post-transcriptional control in ESCs is much less understood. Several lines of evidence indicate that translational regulation provides another important level of control. For example, during mouse ESC differentiation, more than 50% of changes in nuclear protein expression occur without corresponding changes in mRNA levels (1). These data highlight the importance of translational regulation during ESC differentiation and suggest that translational regulators may be key to embryonic development and cell fate determination.

The *Drosophila* Pumilio (Pum), the founding member of the PUF (for <u>Pu</u>milio and <u>F</u>BF [*fem-3 binding factor*]) protein family, has been well-characterized as a translational repressor that directly binds to its target mRNAs. The Pumilio-Homology Domain (Pum-HD), consisting of eight tandem imperfect repeats of 36 amino acids in the C-terminal region of the protein, forms a curved structure in which each repeat contacts one RNA base within the PUF Response Element (PRE), a conserved 32 nucleotide sequence in the 3’ untranslated region (UTR) of Pum target mRNAs (2–5). Upon binding, Pum proteins usually repress translation through both poly(A)-dependent (6–8) and poly(A)-independent pathways (9).

PUF proteins have conserved functions in stem cell proliferation and self-renewal in invertebrates and plants (10, 11). Pum was first identified as a maternal effect mutant required for embryonic patterning in *Drosophila* (12, 13), and has since been implicated in diverse biological processes. In the *Drosophila* ovary, loss of Pum function results in symmetric, rather than asymmetric division of germline stem cells that leads to the depletion of the functional stem cell pool (14–16). The *C. elegans* Pum homologue FBF promotes germline stem cell proliferation and inhibits differentiation (17). PufA, a Pum homologue in *Dyctiostelium*, sustains growth and inhibits differentiation (18), while in Planaria, knockdown of *DjPum* dramatically reduces the number of totipotent stem cells (19). Even in plants, Pum homologues are involved in the regulation of mRNAs involved in shoot stem cell maintenance (20).

Although PUF proteins are highly evolutionarily conserved (21), our understanding of the function of mammalian PUF proteins remains limited. The mouse genome encodes two PUF proteins, Pumilio1 (Pum1) and Pumilio2 (Pum2). *Pum2* mutant ESCs were reported to have no obvious defects in self-renewal or differentiation (22). Correspondingly, *Pum2* mutant mice were viable and fertile with only smaller testes (23) and subtle neurological defects in memory, nesting behavior, and as well as increased propensity to seizures (22, 24). Similarly, little is known about the function of Pum1; conditional knockout of Pum1 in the mouse testis results in decreased fertility (25). Recently, *Pum1-Pum2* double mutant been shown to be embryonic lethal (26) and that Pum1 and Pum2 deficiencies reduces body size in a dose-dependent manner, partly due to translational de-repression of the cell cycle inhibitor CDKN1B (24). Conditional knockout of Pum1 and Pum2 in the nerve system affects neural stem cell maintenance by de-repressing the translation of many mRNAs (27). Pum1 has also been implicated in human neurodegeneration and motor ataxia (28) and the maintenance of genomic stability (29). Despite the well-recognized importance of Pum1 and Pum2 in development, the function of Pum1 and Pum2 proteins in ESCs, either individually or collectively, have not been systematically studied. At the molecular level, Pum1 and Pum2 mRNA targets in ESCs are not known, and the mechanisms whereby Pum proteins regulate gene expression other than translational repression have not been reported.

Here we report the contrasting but complementary functions of *Pum1* and *Pum2* in ESCs that are essential for mouse ESC pluripotency and embryogenesis. We provide evidence that Pum1 and Pum2 form an auto-regulatory, inter-regulatory loop to coordinately control the expression of at least 1,486 mRNAs involved in diverse cellular processes. Furthermore, we demonstrate that such control is not only via translation repression—the well-known mode of Pum regulation--but also by promoting translation and enhancing or reducing stability of different target mRNAs. These findings reveal distinct roles of individual Pum proteins in higher eukaryotes and the novel mechanisms of Pum-mediated regulation of gene expression.

## RESULTS

### Pum1 and Pum2 are expressed in mouse embryonic stem cells and early mouse embryos

To explore Pum function in ESCs and embryos, we first examined the expression of Pum1 and Pum2 in ESCs and embryos by immunofluorescence staining. Pum1 and Pum2 both show diffusely cytoplasmic distributions in interphase and mitotic ESCs and are also present in the nucleus at low levels (Figure 1A, 1B). In metaphase cells, Pum1 and Pum2 appear to be particularly abundant around the nuclear periphery (Fig 1A; arrow). In e2.5 morulae and e3.5 blastocysts, Pum1 and Pum2 are expressed in the cytoplasm (Figure 1C, 1D), with the highest expression in the inner cell mass (ICM) cells of blastocysts and lower expression levels in trophoblast (TB) cells. In e8.5 and e9.5 embryos, Pum1 and Pum2 are primarily expressed in the midbrain, developing somites, and the tail bud, as revealed by whole-mount in situ RNA hybridization (Figure 1E, 1F). The presence of Pum1 and Pum2 in ESCs and early embryos indicate their possible function in ESCs and embryogenesis.

**Figure 1.**
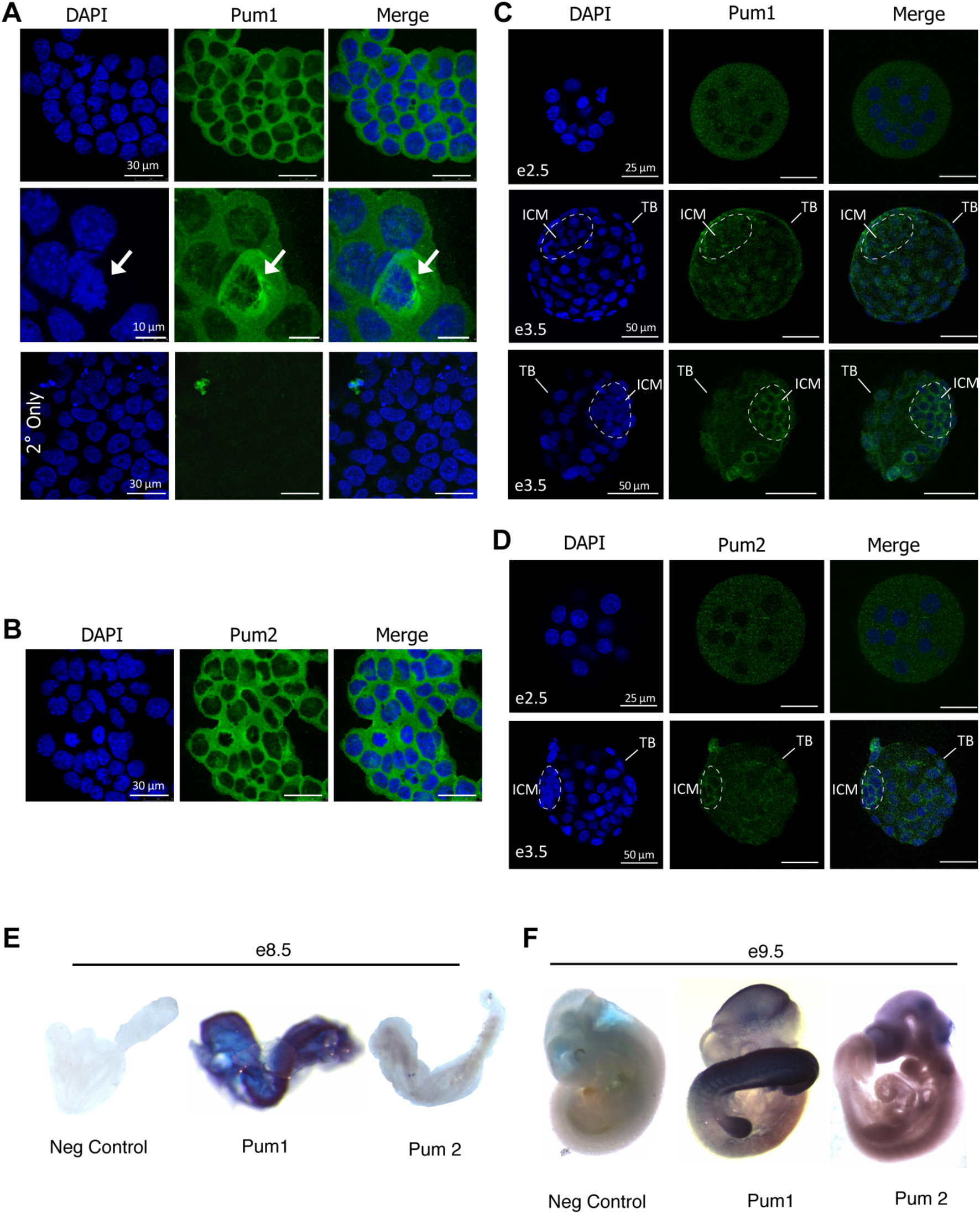
Pum1 and Pum2 expression in ESCs and during early embryogenesis. **(A)** Immunofluorescence microscopic images of Pum1 in CCE wild type ESCs. Pum1 is diffusely cytoplasmic and is enriched in the nuclear periphery of some mitotic cells (arrowhead); minimal background signal is obtained with no primary antibody (lower panel). Nuclei were counterstained with DAPI (blue) in A, B, C, and D. **(B)** Immunofluorescence microscopy of Pum2 in CCE wild type ESCs. Expression is diffusely cytoplasmic. **(C, D)** Immunofluorescence staining of e2.5 morulas (upper panel) and e3.5 blastocysts (lower panels). Pum1 and Pum2 are expressed in both inner cell mass (ICM) and trophoblast (TB) cells, with greater cytoplasmic expression levels in the ICM. **(E, F)** Whole mount *in situ* hybridization analysis of the expression of *Pum1* and *Pum2* was performed on e8.5 (E) and e9.5 (F) wild type embryos; signal is particularly enriched in the forebrain, developing somites, and tail bud.

### *Pum1^−/−^* mice are compromised in viability, prenatal development, and postnatal growth

To determine the function of Pum1 in embryogenesis definitively, we generated *Pum1^−/−^* mice for phenotypic analysis by breeding *Pum1^Flox/+^* mice with *EIIa-Cre* mice using a previously described scheme ((27), Supplemental Figure 1A). Knockout of Pum1 was confirmed by genotyping (Supplemental Figure 1B), quantitative RT-PCR, and western blot analysis, which could not detect either the mutant *Pum1* mRNA nor a truncated protein. The *Pum1^−/−^* mice were much less viable than the expected Mendelian ratio (13% vs 25%, n=244, Supplemental Figure 1C), indicating that loss of *Pum1* compromised prenatal development. In addition, both male and female *Pum1^−/−^* mice were smaller than their heterozygous and wild type littermates at all time points observed (Figure 2A-D), with an average weight 35% less than wild type siblings at 28 dpp. Moreover, aged *Pum1^−/−^* mice had very little body fat, developed a prominent hunchback (Fig 2B), and weighed 43% less than wild type littermates by 11 months. Liver, lung, stomach, intestine, testis, uterus, and brain weights were proportionately smaller (Supplemental Figure 1D), while the testis, kidney, spleen, and heart were disproportionately smaller than in *Pum1^+/−^* or *Pum1^+/+^* littermates (Supplemental Figure 1E). Notably, almost 80% of *Pum1^−/−^* mice were afflicted with ulcerative dermatitis at age 24 weeks, in contrast to 5% of *Pum1^−/+^* littermates (p<0.5; Fig 2E). Histological analysis revealed the most obvious defects are in the testis as reported (25) and in the intestine in which *Pum1^−/−^* mice had blunted, disorganized small intestinal villi (Figure 2F; arrows), but no significant defect in cell proliferation (Ki67 staining, Figure 2F; middle panel), villus length, crypt length, or villus-to-crypt ratio (Figure 2F; lower panel). These findings are consistent with the broad expression of *Pum1* in adult tissues (28) and indicate an important role of Pum1 in postnatal growth.

**Figure 2.**
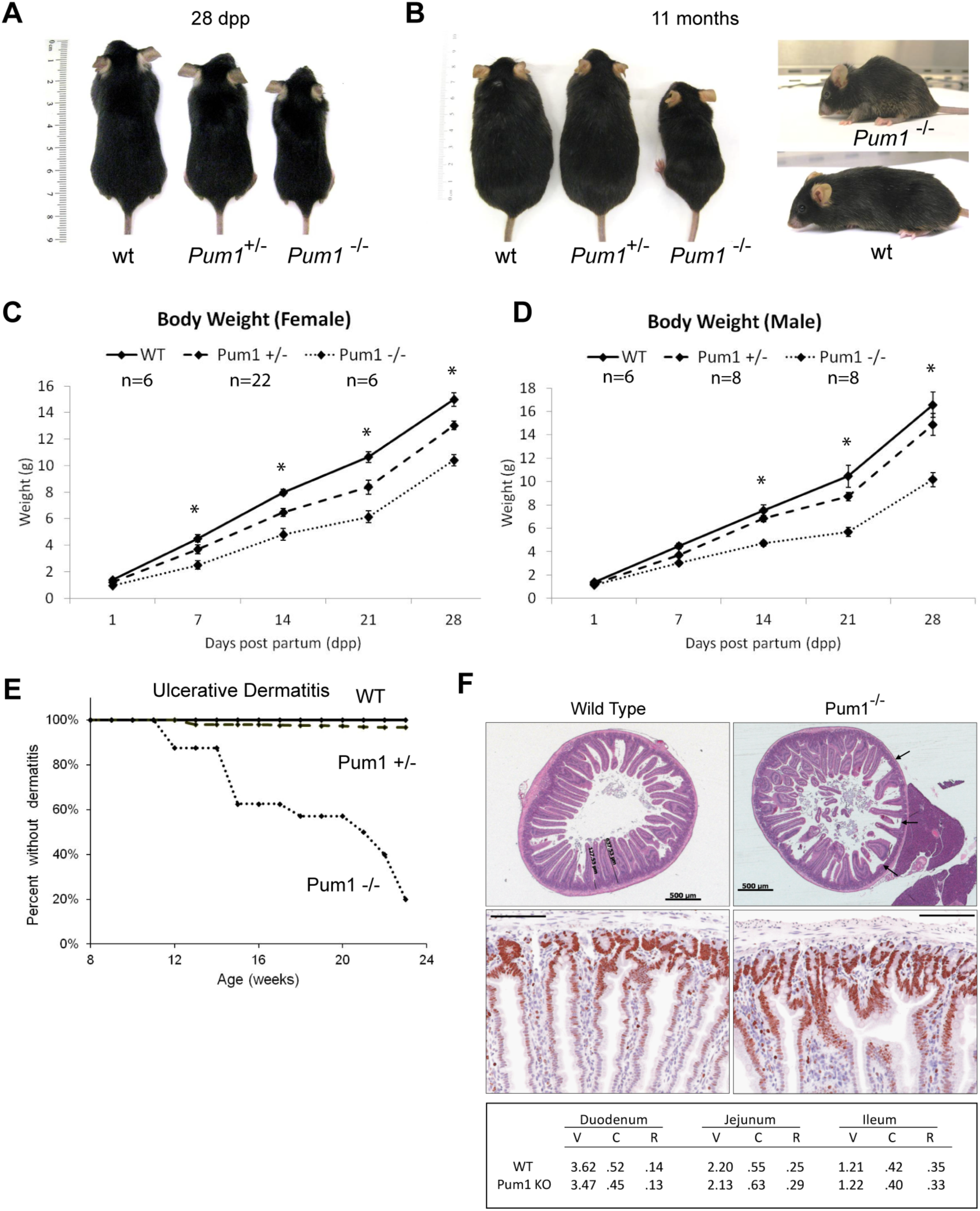
*Pum1* knockout mice have decreased body weight and intestinal defects. **(A)** Phenotype of wild type (wt), *Pum1^+/−^*, and *Pum1^−/−^* littermates at 28 days post partum (dpp). **(B, C)** Phenotype of wild type (wt), *Pum1^+/−^*, and *Pum1^−/−^* littermates at 11 months; *Pum1^−/−^* mice have a hunched appearance (upper right panel) that becomes more prominent with age, and weigh 43% less than wild type mice. (**C-D**) Body weight (in grams) of female (**C**) and male (**D**) wild type (solid line), *Pum1^+/−^* (dashed line), and *Pum1^−/−^* (dotted line) littermates at 1, 7, 14, 21, and 28 days post partum (dpp), with bars indicating SEM. * indicates p value < 0.01. At 28 dpp, *Pum1^−/−^* mice weigh 35% less than wild type mice. **(E)** Kaplan-Meier plot of the relative risk of ulcerative dermatitis in wild type (solid line), *Pum1^+/−^* (dashed line), and *Pum1^−/−^* (dotted line) in 8-24 week old mice. At 11 weeks, 16% of *Pum1^−/−^* mice have ulcerative dermatitis compared to 0% of wild type mice, and at 23 weeks, 80% of *Pum1^−/−^* mice have ulcerative dermatitis compared to 5% of wild type littermates. **(F)** Histological analysis of *Pum1* −/− mice. Thin sections of the duodenum of wild type and Pum1 knockout mice were stained with H & E (upper panel); multiple areas within the plane of sectioning (arrows) are representative of blunted, disorganized villi in the duodenum of Pum1 −/− mice. Ki67 staining (middle panel) is comparable in wild type (left) and Pum1 −/− (right) duodenum sections. Villus vs. crypt length (lower panel) is not statistically different between wild type and Pum1 −/− mice in the duodenum, jejnum, or ileum. V: villus, C: crypt, R: villus/crypt ratio.

### The function of Pum1 and Pum2 are partially redundant and dose-sensitive

The relatively mild phenotype of *Pum1^−/−^* mice could be due to the high degree of homology between Pum1 and Pum2 (91% identity and 97% similarity in the Pum-HD domain) (30), which renders them functionally redundant. To address this possibility, we generated *Pum2^+/−^* mice (Supplemental Figure 2A, 2B). These mice showed normal viability and body weight at all time points assessed (Supplemental Figure 2D-F).

To determine whether Pum1 and Pum2 could have redundant or dose-dependent functions, *Pum1^+/−^; Pum2^+/−^* mice were crossed to obtain nine possible genotypes ranging from zero to four functional *Pum* alleles. At 1dpp, all genotypes were recovered except for *Pum1^−/−^; Pum2^−/−^* mice, indicating that these embryos do not survive to birth (Figure 3A; n=127, p value <0.01). In addition, *Pum1^−/−^; Pum2^+/−^* pups were recovered at significantly lower numbers than expected (Figure 3A; p<0.01) and were smaller than their littermates at 1dpp, had no milk in their stomach, and mostly died within 24 hours of birth (Figure 3B), with only one surviving to 6dpp. Histological analysis of *Pum1^−/−^; Pum2^+/−^* pups showed thymic necrosis, hepatic congestion, and hepatic atrophy (data not shown), but no obvious defects in heart, lung, kidney, spleen, or stomach. In contrast, *Pum1^+/−^; Pum2^−/−^* mice were relatively normal, whereas *Pum1^−/−^* mice were smaller than *Pum2^−/−^* mice, which were in turn smaller than *Pum1^+/−^; Pum2^+/−^* mice (Figure 3C). In fact, this trend is already evident during embryogenesis at e9.5 and e12.5 (Supplemental Results and Supplemental Figure 3). These observations support a partially redundant and collectively essential role of Pum1 and Pum2 during embryogenesis, with Pum1 having greater effects than Pum2.

**Figure 3.**
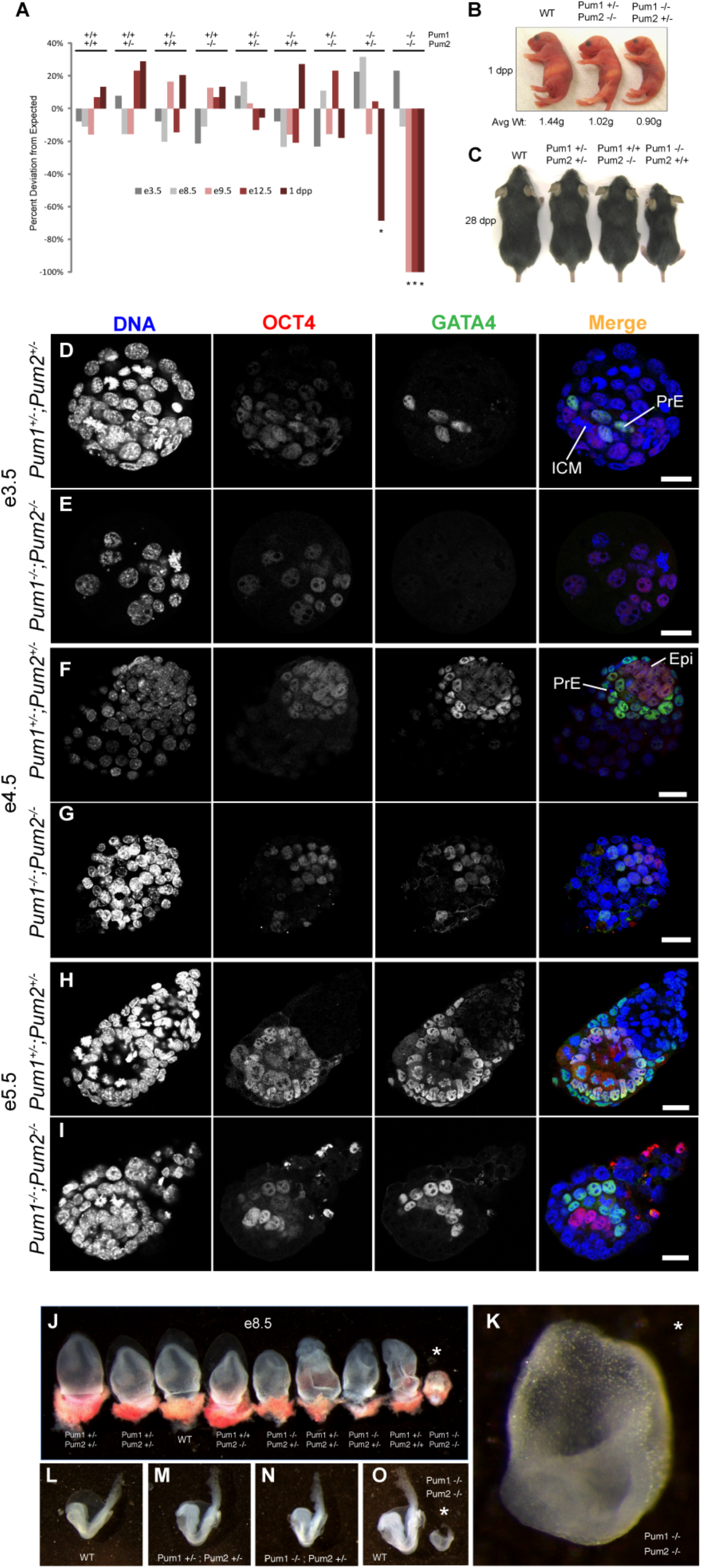
*Pum1^−/−^; Pum2^−/−^* mice are embryonic lethal by e8.5. **(A)** Deviation of the ratio of observed from expected genotypes of the *Pum1^+/−^; Pum2^+/−^* x *Pum1^+/−^; Pum2^+/−^* mating at e3.5 (dark grey, n = 52), e8.5 (light gray, n=36), e9.5 (pink, n=21), e12.5 (red, n=23), and 1 day post partum (dpp; dark red, n=127). The observed ratio of *Pum1^−/−^; Pum2^+/−^* pups is 65% less than expected at 1dpp, and of *Pum1^−/−^; Pum2^−/−^* embryos is 100% less than expected at e9.5, e12.5, and 1dpp. * and ***indicates p < 0.01 and <0.001, respectively. **(B)** Morphology and average weight (in grams) of wild type, *Pum1^+/−^; Pum2^−/−^*, and *Pum1^−/−^; Pum2^+/−^*, littermates at 1dpp. *Pum1^+/−^; Pum2^−/−^* pups weigh 29% less than WT littermates; *Pum1^−/−^;Pum2^+/−^* pups weigh 38% less than WT littermates, have no milk in their stomachs, and die within 24 hours of birth. **(C)** Phenotype of Pum-deficient mice at 28 dpp. *Pum1^−/−^* mice are smaller than *Pum2^−/−^* mice and *Pum1^+/−^; Pum2^+/−^* mice, which are both indistinguishable from wild type at 28dpp. **(D-I)** *pum1^−/−^; pum2^−/−^* embryos are developmentally delayed from e3.5dpc-e5.5. (D, E) Wildtype and mutant embryos at e3.5 were stained for Oct4 (red) and Gata4 (green). No or smaller blastocoel cavity exist in the mutant embryos. Furthermore, Gata4-positive primitive endoderm (PrE) cells were absent in the mutant embryos. (F, G) Immunofluorescence staining of wildtype (E) and mutant (G) embryos at 4.5dpc. PrE cells are randomly positioned in the inner cell mess (ICM) in mutant embryos. (H, I) Wildtype (H) and mutant (I) embryos at 5.5dpc were stained for Oct4 and Gata4. Wildtype embryos developed an inner layer of epiblast cells (Oct4-positive) and an outer layer of visceral endoderm cells (Gata4-positive). However, in mutant embryos, Gata4-positive PrE cells are still next to the blastocoel cavity. Nuclei were stained with DAPI (blue). Scale bars, 25µm. **(J)** Embryonic and extraembryonic tissues dissected from uteri of *Pum1^+/−^; Pum2^+/−^* mated mice at e8.5. Yolk sacs were used for genotyping. A *Pum1^−/−^; Pum2^−/−^* embryo (far right) is smaller than all littermates. **(K-O)** Higher magnification of dissected e8.5 embryos; *Pum1^−/−^; Pum2^−/−^* embryo shows developmental delay, primitive head fold, and overall lack of tissue with especially thin neural tissue.

### *Pum1^−/−^; Pum2^−/−^* embryos show developmental delay at the morula stage and are lethal by e8.5

To determine the embryonic defects that lead to the lethality of *Pum1^−/−^; Pum2^−/−^* mice, given the known stem-cell function of Pum proteins in lower organisms, we first examined *Pum1^−/−^; Pum2^−/−^* and sibling *Pum1^+/−^; Pum2^+/−^* e3.5 blastocysts in which the inner cell mass has just been established. These embryos were isolated from uteri and genotyped individually by DNA extraction followed by nested PCR. Four *Pum1^−/−^; Pum2^−/−^* embryos were recovered from 52 blastocysts (Supplemental Figure 4A) at the expected Mendelian ratio, indicating that double knockout embryos are viable up to e3.5. However, two of the four *Pum1^−/−^; Pum2^−/−^* embryos still appeared morula-like without a defined blastocoel cavity (Supplemental Figure 4A, asterisks), whereas all 48 of the remaining e3.5 embryos that carried at least one wildtype Pum1 or Pum2 allele had progressed to the blastocyst stage. To obtain a better quantification of the penetrance of the *Pum1^−/−^; Pum2^−/−^* phenotype, we isolated another 17 double mutant e3.5 embryos from doxycycline-treated *Pum1^flox/flox^, Pum2^flox/flox^; rtTA*[*ROSA*]*26-Cre* mice (see Materials and Methods), of which eight appeared as morulae and only nine developed to blastocysts (Figure 3E). In contrast, among 28 *Pum1^+/−^; Pum2^+/−^* e3.5 embryos, only one appeared morula-like, but all 27 others were blastocysts (Figure 3D). Hence, 10 out of 21 (47%) *Pum1^−/−^; Pum2^−/−^*e3.5 embryos were still at the morula stage, in contrast to 1 out of 75 (1.33%) control e3.5 embryos (Supplemental Table 1). To determine if the 47% *Pum1^−/−^; Pum2^−/−^*morulae were due to developmental delay or arrest, we cultured the them overnight and they all developed into blastocysts (Supplemental Figure 4B), indicating a developmental delay. The incomplete penetrance of the delayed phenotype further indicates that the defects of *Pum1^−/−^; Pum2^−/−^* embryos might have just begun to manifest at this stage.

To further investigate the nature of the lethality of *Pum1^−/−^; Pum2^−/−^* embryos, we examined the morphology and expression of key cell-fate markers in doxycycline-induced *Pum1^−/−^; Pum2^−/−^* embryos at e3.5, e4.5, e5.5, e6.5, e7.5, e9.5, and e12.5 stages and *Pum1^+/−^; Pum2^+/−^* control embryos at the same stages (see Materials and Methods). At 3.5dpc, the *Pum1^−/−^; Pum2^−/−^* morulae express the pluripotency marker Oct4 normally but do not express the endoderm marker Gata4 (Figure 3E). This indicates that *Pum1^−/−^; Pum2^−/−^* embryos already show a defect in endoderm lineage differentiation at e3.5.

During subsequent embryogenesis, the *Pum1^−/−^, Pum2^−/−^* embryos further display defects in differentiation. By e4.5, 56% of the mutant embryos showed developmental delay or arrest (Supplementary Table 1). In these embryos, the Gata4-positive primitive endoderm cells are randomly positioned in the inner cell mass, equivalent to 3.5dpc wild type embryos and indicating a delay in this lineage (Figure 3G, Supplemental Figure 4C-D). By e5.5, 62% of the mutant embryos show incomplete epiblast cell transformation, with Gata4-positive cells still positioned next to the blastocoel cavity (*cf.* Figure 3H, I). This mislocalization continues in most of the 6.5dpc *Pum* mutant embryos (*cf.* Supplemental Figure 6 A, B). The trophectoderm lineage, as indicated by Cdx2 expression, developed normally from e3.5 to e5.5 in the mutant embryos (*cf.* Supplemental Fig. 5), but by e6.5, Cdx2-positve cells are no longer detectable (Supplementary Figure 6B). By 7.5dpc, all cells in the mutant embryos become disorganized, and no obvious structure is observable (*cf.* Supplementary Figure 6C, D). By e8.5, only two *Pum1^−/−^; Pum2^−/−^* embryos were recovered from 36 embryos derived from a *Pum1^+/−^; Pum2^+/−^* self-cross; both were significantly smaller and ill-developed (Figure 3J). They have a primitive head fold but overall lack of tissue with thin neural tissue (Figure 3 K-O). No *Pum1^−/−^; Pum2^−/−^* embryos were recovered at e9.5 or e12.5 (Figure 3A). Therefore, the major defects observed at e7.5 likely reflect the terminal phenotype immediately before the e8.5 lethal phase of the mutant embryos.

### *Pum1, Pum2*, and *Pum1;Pum2* deficiencies do not significantly affect the proliferation or viability of ESCs

Because *Pum1^−/−^; Pum2^−/−^* embryos display developmental delay at the morula-to-blastocyst transition at which ESCs form, we further investigated the function and mechanisms of action of Pum1 and Pum2 in ESCs. ESCs also allowed us to bypass the challenge of studying such mechanisms in scarcely available *Pum* double mutant embryos at this stage. We first derived *Pum1^Flox/+^* and *Pum1^Flox/Flox^* ESC lines from e3.5 blastocysts of *Pum1^Flox/+^* mice. The ESC lines were transfected with a *pBabe-Puro-Cre* plasmid expressing Cre recombinase, and selected in media containing puromycin to generate wild type, *Pum1^+/−^*, and *Pum1^−/−^* cell lines, respectively. Knockout of *Pum1* was confirmed by genotyping, quantitative RT-PCR, and western blot analysis (Supplemental Figure 7A-C), which indicate that neither the mutant mRNA nor a truncated protein is detectably expressed.

However, repeated attempts to derive a *Pum1^−/−^; Pum2^−/−^* ESC line were unsuccessful, either from blastocysts of *Pum1^+/−^; Pum2^+/−^* offspring or from Cre-mediated excision of *Pum1^Flox/Flox^;Pum2^Flox/Flox^* ESCs. Hence, we transfected *Pum1^Flox/Flox^;Pum2^Flox/Flox^* cell lines carrying an inducible doxycycline-responsive Cre gene (rtTA[ROSA]26) with a doxycycline-inducible *Pum1* cDNA construct. Addition of doxycycline-induced deletions in both *Pum1* and *Pum2* genes while concurrently inducing the expression of the exogenous *Pum1* cDNA to rescue the *Pum1-and-Pum2* deficiency (see Materials and Method). The genotype of the resulting *Pum^−/−^; Pum^−/−^* ESCs was confirmed by western blotting (Supplemental Figure 7D; I). These ESCs were maintainable after the withdrawal of doxycycline to stop *Pum1* expression. Their proliferation was then compared to that of WT, *Pum1^−/−^*, and *Pum2^−/−^* ESCs when cultured with or without feeders. The cell doubling times of all of the ESCs cell lines are approximately 15 hours (Supplemental Figure 7E-H). More double mutant ESCs are positive for Annexin V, an early apoptotic marker and for propidium iodide, a marker for pyknotic nuclei than wildtype ESCs, but this difference is not statistically significant (Supplemental Figure 7I-J), indicating that Pum1 and Pum2 are not essential for the viability of ESCs.

### Pum1 promotes ESC differentiation whereas Pum2 promotes ESC self-renewal; together they are required for ESC pluripotency

To pinpoint the defects of *Pum1^−/−^*, *Pum2^−/−^* and *Pum1^-/,^*;*Pum2^−/−^* ESCs, we cultured wild type, *Pum1^−/−^, Pum2 ^−/−^*, and *Pum1^−/−^;Pum2 ^−/−^* ESCs under a spectrum of differentiation-promoting conditions. Under normal culture conditions with feeder cells in the presence of LIF, the loss of either *Pum1* or *Pum2* did not affect the percentage of AP–positive colonies, but the loss of both *Pum1* and *Pum2* caused a more significant decrease in AP–positive colonies (Figure 4A-C), indicating the decreased self-renewal capacity of *Pum1^−/−^*; *Pum2^−/−^* ESCs. However, under a mild differentiation-promoting culture condition without feeder cells, *Pum1^−/−^* cells generated more AP-positive colonies and *Pum2^−/−^* ESCs generated fewer AP-positive colonies than wild type ESCs, while the double knockout ESCs generated an intermediate number of AP-positive colonies (Figure 4A, B, D). These results indicate that, under mild differentiation-promoting conditions, Pum1 functions to promote differentiation as previously reported for haploids ESCs (31) whereas Pum2 functions to promote self-renewal, so the overall effect of losing both genes yielded an intermediate result.

**Figure 4.**
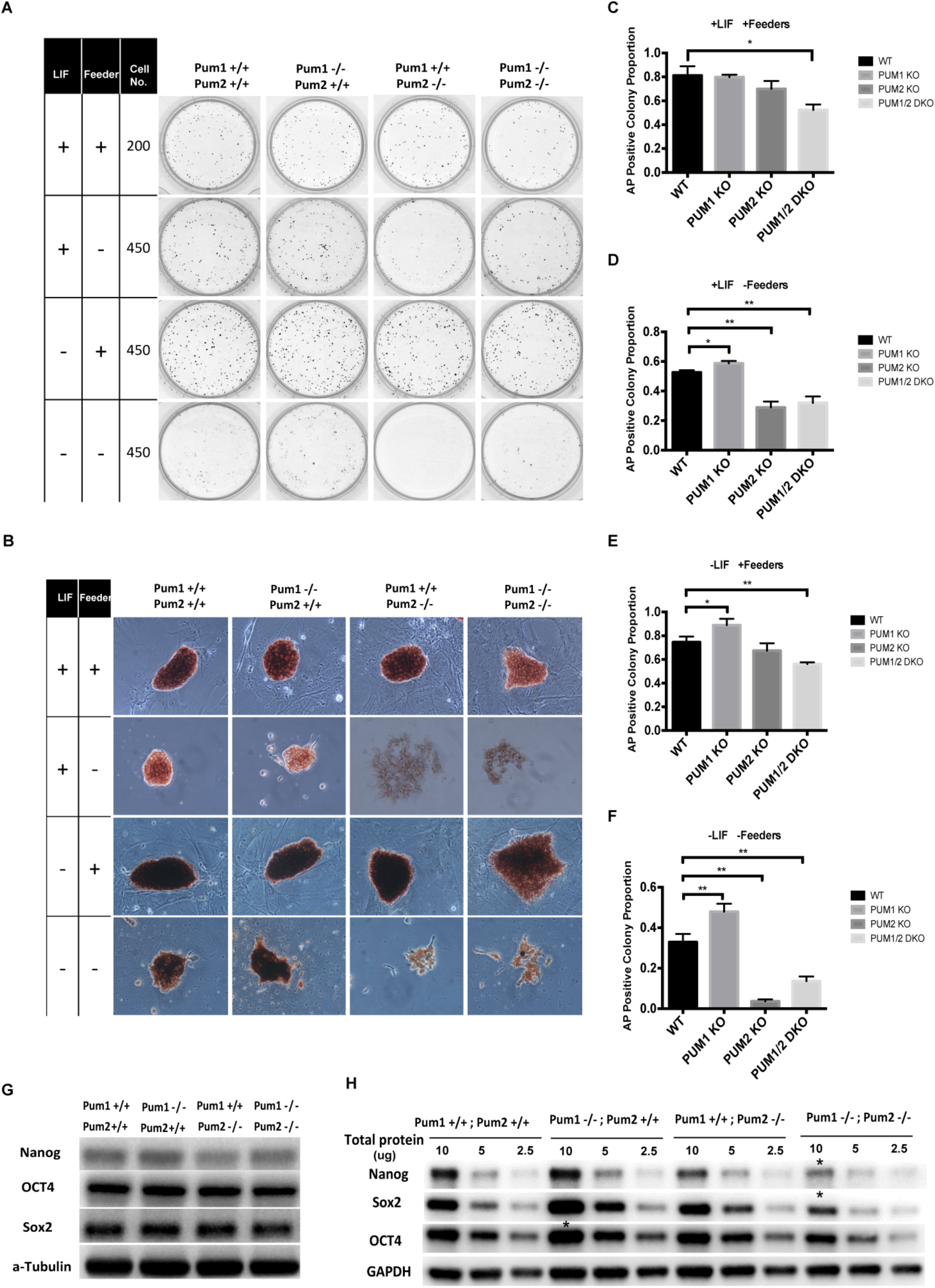
*Pum1;Pum2* double knock-out (DKO) ESCs showed decreased self-renewal capacity. (**A, B**) Alkaline phosphatase (AP) staining of ESCs cultured under self-renewal and differentiation conditions as viewed by low (A) and high (B) magnifications. (**C-F**) Quantification of AP staining in different culture conditions. Cell colony numbers are shown as mean±SD. T-test, *p<0.05, ** p<0.01. (**G, H**) Western blot analysis of pluripotent markers in wildtype, *Pum1^−/−^, Pum2^−/−^*, and *Pum1^−/−^; Pum2^−/−^* ESCs cultured under self-renewal (G) and differentiation (H) conditions. Asterisks in H highlight lower protein levels of Nanog and Sox2 in *Pum1^−/−^; Pum2^−/−^* mESCs and higher protein levels of Oct4 in *Pum1^−/−^* mESCs compared to wild type.

To confirm this, we cultured ESCs under another moderate differentiation-promoting condition with feeder cells but without LIF. We again observed increased number of AP-positive colonies from *Pum1^−/−^* ESCs and decreased number of AP-positive colonies from *Pum2^−/−^* ESCs (Fig 4A, B, E), which supports the above conclusion. However, the double mutant ESCs generated somewhat fewer AP-positive colonies. Given that feeders cells not only support pluripotency but also cell survival, this could reflect that the double mutant ESCs are more dependent on the survival-promoting function of feeder cells. We then removed both LIF and feeder cells. Under this highly differentiation-promoting condition, *Pum1^−/−^* ESCs again generated more AP-positive colonies whereas *Pum2^−/−^* ESCs generated fewer AP-positive colonies than wild type ESCs. The double mutant ESCs again generated an intermediate number of AP-positive colonies (Figure 4A, B, F). Overall, these data support a model in which Pum1 promotes ESC differentiation and Pum2 promotes ESC self-renewal.

The above conclusion is also supported by expression levels of key pluripotency genes in *Pum1^−/−^*, *Pum2^−/−^*, and double mutant ESCs. When cultured with feeder cells and LIF, *Pum1^−/−^*, *Pum2^−/−^*, and double mutant ESCs all expressed similar levels of Nanog, Sox2, and Oct4 as wild type ESCs (Figure 4G). However, without feeder cells and LIF, *Pum1^−/−^* ESCs expressed higher levels of Nanog, Sox2, and Oct4 than wild type ESCs; whereas *Pum2^−/−^* ESCs expressed a lower level of Nanog than wild type ESCs (Figure 4H). Double knockout ESCs showed significantly reduced expression of Nanog and Sox2, and somewhat reduced expression of Oct4 (Figure 4H). These results confirm that Pum1 promotes differentiation whereas Pum2 promotes self-renewal, so that losing both genes severely compromises ESC pluripotency.

### *Pum2^−/−^* ESCs show precocious expression of differentiation genes yet *Pum1^−/−^*;*Pum2^−/−^* ESCs are severely defective in differentiating into three germ layer lineages

To further investigate the role of Pum1 and Pum2 in ESC self-renewal and differentiation, embryoid bodies (EBs) were generated from *Pum1^−/−^*, *Pum2^−/−^*, and double mutant ESCs and assayed for markers of pluripotency as well as endodermal, mesodermal, and ectodermal lineages (Figure 5A-J). *Pum1^−/−^* and *Pum2^−/−^* EBs showed a similar rate of growth and morphology as wild type EBs up to 20 days. However, beating cardiomyocytes appeared earlier in *Pum2^−/−^* EBs (16 days) compared to wild type and *Pum1^−/−^* EBs (20 days), indicating an accelerated differentiation of *Pum2^−/−^* ESCs along the mesodermal lineage. In support of this notion, *Pum2^−/−^* EBs showed precocious expression of mesodermal markers Brachyury and Goosecoid, both peaking at Day 4 (Figure 5C, D). Moreover, Pum2^−/−^ EBs had accelerated expression of ectodermal marker FGF5 and endodermal markers Foxa2, Gata4 and Gata6. Together, the above observations indicate role of Pum2 in promoting self-renewal and pluripotency.

**Figure 5.**
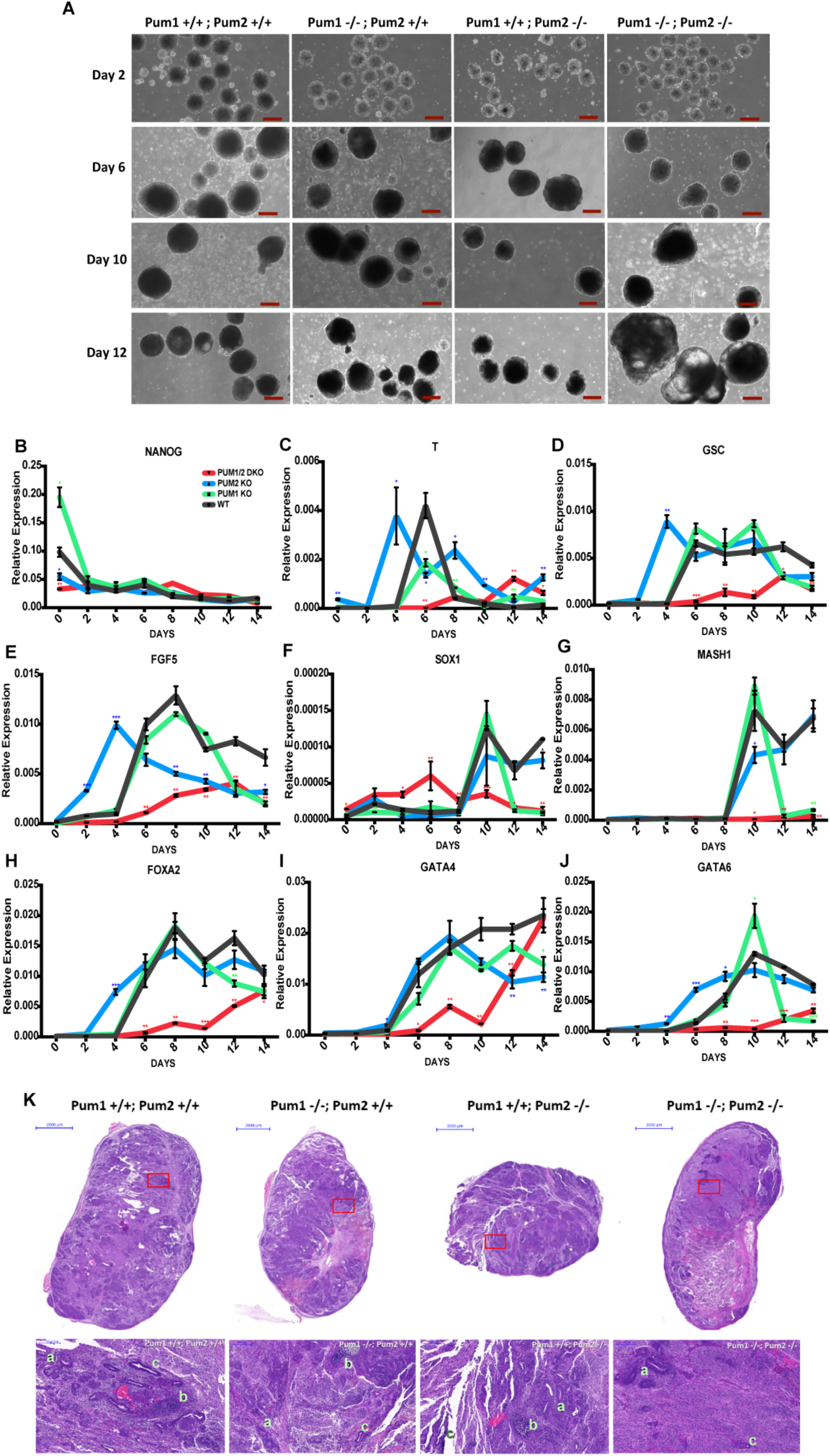
*Pum1;Pum2* DKO ESCs are severely defective in expressing key marker of the three germ layers in early differentiation simulated by the embryoid body and teratoma assays. **(A)** Morphological Comparison between wildtype and mutant EBs in Day 2, Day 6, Day 10 and Day 12. (**B-J**) Relative expression of markers for pluripotency and three germ layers as quantified by qRT-PCR and normalized over *GAPDH* mRNA. Results are presented as mean ± SD. T-test, *p<0.05, ** p<0.01, *** p<0.001. (**K**) H&E staining of teratoma histological sections. a. ectoderm-like pattern; b. mesoderm-like pattern; c. endoderm-like pattern.

In contrast, Pum1/2 double mutant EBs initially grew much slower but eventually reached wild type size by Day 8-10, and became much larger than wild type EBs by Day 12 (Figure 5A). They formed prominent cysts indicative of primitive ectoderm differentiation (Fig 5A), but had no beating cardiomyocytes even beyond Day 20, consistent with a defect in mesodermal differentiation. The expression of the pluripotency marker Nanog expectedly decreased over the 14-day course of differentiation (Figure 5B), but the expression of mesodermal markers Brachyury and Goosecoid, ectodermal markers FGF5, Sox1 and Mash1, and endodermal marker Gata6 all failed to increase even after 14 days in culture (Figure 5C-G, J). The expression of two other endodermal markers Foxa2 and Gata4 were also severely delayed up to Day 10 (Figure 4H, I). These observations indicate that *Pum1^−/−^;Pum2^−/−^* ESCs are severely defective in differentiation, in addition to impaired self-renewal as described above.

### *Pum1^−/−^*; *Pum2^−/−^* ESCs do not properly differentiate into three germ layer lineages in teratomas

To confirm the *in vitro* differentiation defects of Pum-deficient ESCs, four cell lines were injected subcutaneously for teratoma assays (Figure 5K, Supplemental Figure 8). The growth rate of *Pum1^−/−^;Pum2^−/−^* teratomas was transiently higher than that of *Pum1^−/−^*, *Pum2^−/−^*, and wild type cells, but eventually showed no significant difference (Supplemental Figure 8A-C). However, hematoxylin-eosin staining revealed that the double mutant teratomas contained many more undifferentiated cells and lacked mesoderm-like structures (Figure 5K, for typical mesodermal structures, see Supplemental Figure 8D). In addition, the double mutant teratomas contained fewer Sox1-positive ectoderm structure and Foxa2-positive endodermal cells (Supplemental Figure 8E, F). Together, these data confirm that double mutant ESCs are defective in *in vivo* differentiation into the three germ lineages.

### Pum1 and Pum2 form an auto- and inter-regulatory feedback loop in ESCs

To further investigate the functional relationship between Pum1 and Pum2, we found multiple putative PRE sequences in the 3’UTR of both Pum1 and Pum2 (Figure 6A). Therefore we reasoned that Pum1 and Pum2 likely repress their own and each other’s expression. To investigate this possibility, we examined the expression of Pum1 in *Pum2^−/−^* ESCs and vice versa. Indeed, the levels of Pum1 and Pum2 are significantly increased in *Pum2^−/−^* and *Pum1^−/−^* backgrounds, respectively, illustrating that Pum 1 and Pum2 repress each other’s expression (Figure 6A). Such a negative auto- and inter-regulatory feedback loop would maintain steady-state levels of Pum1 and Pum2, meanwhile allowing one Pum to be over-expressed in the absence of the other, which may partially compensate for defects caused by the loss of either individual protein.

**Figure 6.**
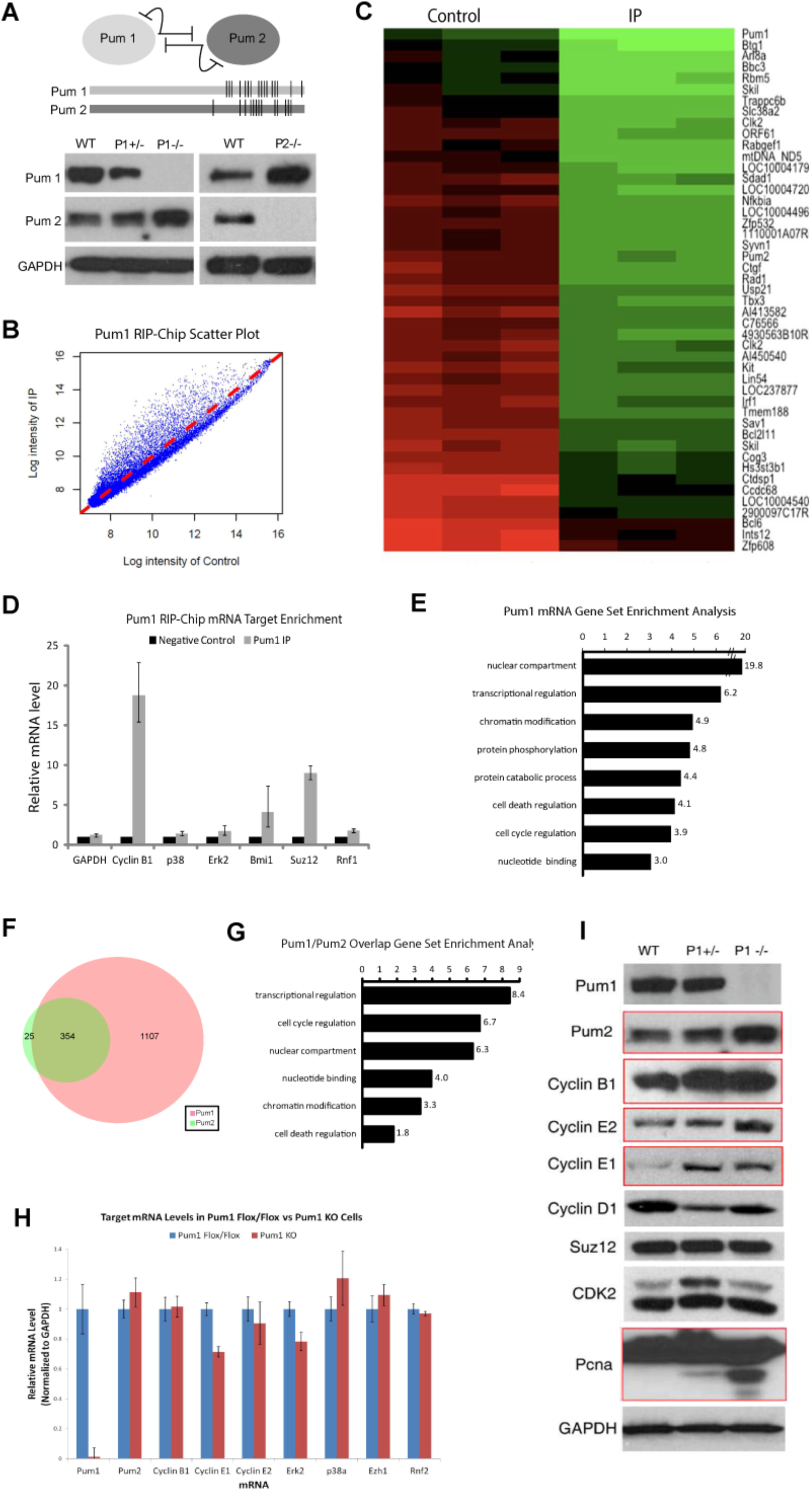
Pum1 and Pum2 auto-regulate each other and bind overlapping sets of functionally related mRNAs in mESCs. **(A)** Pum1 and Pum2 participate in an autoregulatory loop (upper panel); *Pum1* and *Pum2* mRNA have multiple putative PRE sites in their 3’ UTR (middle panel); Western blotting analysis reveals increased Pum2 levels in *Pum1^−/−^* ESCs (lower left panel) and increased Pum1 levels in *Pum2^−/−^* ESCs (lower right panel). **(B)** Scatter plot of the log intensity of IP samples (y axis) and of negative controls (incubated with blocking peptide against which the antibody was raised [x axis]). **(C)** Heat map of enrichment in IP and control samples for the top 50 genes. **(D)** qRT-PCR of known and novel Pum1 mRNA targets confirming enrichment in IP samples; error bars indicate SEM of three replicates. **(E)** Gene Set Enrichment Analysis (GSEA) of top 500 Pum1 mRNA targets by fold enrichment and *p* value. **(F)** Venn diagram indicating total number of significantly enriched (p<0.001, fold enrichment >1.5) mRNA targets of Pum1 (pink) and Pum2 (green). **(G)** GSEA of the 354 overlapping mRNA targets of Pum1 and Pum2. **(H)** qRT-PCR of Pum-1 mRNA targets identified by RIP-Chip in Pum1 floxed (blue) and *Pum1* knockout (red) cells. Error bars represent SEM from 3 replicates. **(I)** Western blot analysis of Pum1 mRNA targets in wild type, *Pum1^+/−^*, and *Pum1^−/−^* ESC lines indicating translational upregulation of several targets in *Pum1^−/−^* cells.

### Pum1 binds to the mRNAs of 1461 genes in ESCs

Given the well-established role of Pumilio proteins as translational repressors, we next sought to identify mRNA targets of Pum1 and Pum2 in ESCs to explore potential mechanisms of the observed defects in ESC self-renewal and differentiation. Endogenous Pum1 immunoprecipitation-microarray (RIP-Chip) experiments were conducted as previously described (32) on total ESC lysates to identify the associated mRNAs (Supplemental Figure 9A,B). Data were normalized using loess regression and quantile normalization; the p values and fold enrichment for each probe were analyzed by a Volcano plot and heat maps of probe enrichment (Figure 6B, C, Supplemental Figure 9C-E).

1461 unique mRNAs were significantly enriched in the Pum1 IP as compared to the negative control (> 1.5 fold, p value <0.001) (Figure 6C). Several individual targets including Cyclin B1, Suppressor of Zeste 12 homolog (Suz12), and B lymphoma Mo-MLV insertion region 1 homolog (Bmi1) were verified by RIP and quantitative RT-PCR (Figure 6D). Of the 1461 genes, the top 500 by *p* value and fold enrichment (Figure 6C, Appendix A) were assessed by Gene Set Enrichment Analysis (GSEA) to assign the enriched targets to functionally related gene sets. GSEA revealed that Pum1 mRNA targets are particularly enriched for genes involved in transcriptional regulation (23%), cell growth, maturation and death (20%), protein phosphorylation (13%), GTPase activity (9%), embryonic development (8%), ubiquitination (6%), cell cycle regulation (3%), and chromatin regulation (3%) (Figure 6E). To examine whether the targets identified from our RIP-ChIP data display the known Pum binding motif termed Pumilio Response Element (PRE, TGTA[ATC]ATA), we searched for this motif at their 3’-UTR sequences (25, 33) by PERL regular expression search. We found that 52% of Pum1-target genes and 65% of Pum2-targeted genes have at least one motif, indicating that PRE is important but not essential for Pum targeting.

### Pum2 binds to the mRNAs of 379 genes in ESCs

The high homology between Pum1 and Pum2 (34) implies that they may have similar or overlapping target sets. To investigate this possibility, we performed a Pum2 RIP-Chip to identify its *in vitro* mRNA targets, using the same experimental conditions and data normalization as Pum1-RIP-Chips. Gene set enrichment analysis of the top mRNA targets revealed that, like Pum1, Pum2 binds to mRNAs in ESCs that are functionally enriched for proteins involved in transcriptional control, cell cycle regulation, and metabolism. Further analysis revealed that of the 379 most enriched mRNA targets of Pum2 (>1.5 fold, p value <0.001) (Appendix B), 354 were also Pum1 targets that were enriched for proteins involved in transcriptional regulation, cell cycle regulation, and nuclear proteins (Figure 6G), while 25 were Pum2-specific (Figure 6F, G, Appendix C).

### Pum1 represses the translation of Pum2, Cyclin B1, Cyclin E1, and Cyclin E2 mRNAs

To determine whether Pum1 and Pum2 translationally control their identified target mRNAs in ESCs, we compared the steady-state mRNA and protein levels of several targets in wild type, *Pum1^+/−^*, and *Pum1^−/−^* ESC lines. There was no significant difference in the mRNA levels of *Pum2*, *Cyclin B1*, *Cyclin E1*, and *Cyclin E2* (Figure 6H). However, their protein levels were significantly increased by approximately two-fold in Pum1-deficient ESCs (Figure 6I), indicating that Pum1 translationally represses these targets *in vivo*.

To further assess whether the translational repression is mediated by PREs, we utilized a luciferase reporter construct in which the 3’ UTRs of *Cyclin B1* mRNA, which contains three PREs, was cloned directly downstream of the protein coding sequence of firefly luciferase (Supplemental Figure 9F). A plasmid encoding renilla luciferase was co-transfected into cells as a transfection control, and the ratio of firefly to renilla luciferase was used to indicate the effects of Pum proteins on the translation of target mRNAs *in vivo*. The full-length Cyclin B1 3’UTR results in an 80% reduction of firefly luciferase expression levels (Supplemental Figure 9F). We then generated mutations within the eight-nucleotide core of each of the three PREs within the 3’UTR of *Cyclin B1*, as previously reported (35). Mutating each PRE resulted in a 10% to 40% release of the suppression of luciferase expression levels, and mutating all three PREs allowed for a 50% increase of firefly expression. This indicates that Pum proteins suppress Cyclin B1 translation mainly via binding to PREs.

### Pum1 and Pum2 impact the abundance of hundreds of mRNAs in diverse cellular processes

To systematically analyze the effect of Pum1 and Pum2 on mRNAs and their translation in ESCs, we first investigated the changes of the transcriptome in *Pum1^−/−^*, *Pum2^−/−^* and double mutant ESCs by RNA deep sequencing. In *Pum1^−/−^* ESCs, 45 genes are upregulated, yet 8 genes are downregulated (Supplemental Figure 10A, B, G). In *Pum2^−/−^* ESCs, however, 880 genes are upregulated, yet 481 genes are downregulated (Supplemental Figure 10C, D). The up-regulated transcripts are enriched in development, cell cycle, cell proliferation, cell differentiation and apoptotic pathways, whereas the down-regulated gene are enriched in nucleosome assembly and chromatin-related activities (Supplemental Figure 10H, I). In *Pum1^−/−^*;*Pum2^−/−^* ESCs, 773 genes are upregulated, yet 475 genes are downregulated (Supplemental Figure 10E, F). The up-regulated transcripts are also enriched in development, cell cycle, and apoptotic pathways, whereas the down-regulated gene are also enriched in development, nucleosome assembly, and neuron development activities (Supplemental Figure 10J. K).

### Pum1 and Pum2 impact the translation of hundreds of mRNAs in diverse cellular processes

We then systematically analyzed the effect of Pum1 and Pum2 on translation of all mRNAs in ESCs by ribosome protection assays of *Pum1^−/−^*, *Pum2^−/−^* and the double mutant ESCs. We first assessed the quality of our ribosomal protection assay by analyzing the distribution of ribosome footprints on mRNAs at single nucleotide resolution. The ribosome footprints are predominantly 28-nucleotide long (the width of the ribosome; Figure 7A). The footprint shows clear tri-nucleotide periodicity in all samples (Figure 7B, Supplemental Figure 11 A-I), and most footprint reads are in the protein-coding region, but not 5’ or 3’ untranslated regions (UTRs; Figure 7C). Together the above findings validate the high quality of our ribosome protection data.

**Figure 7.**
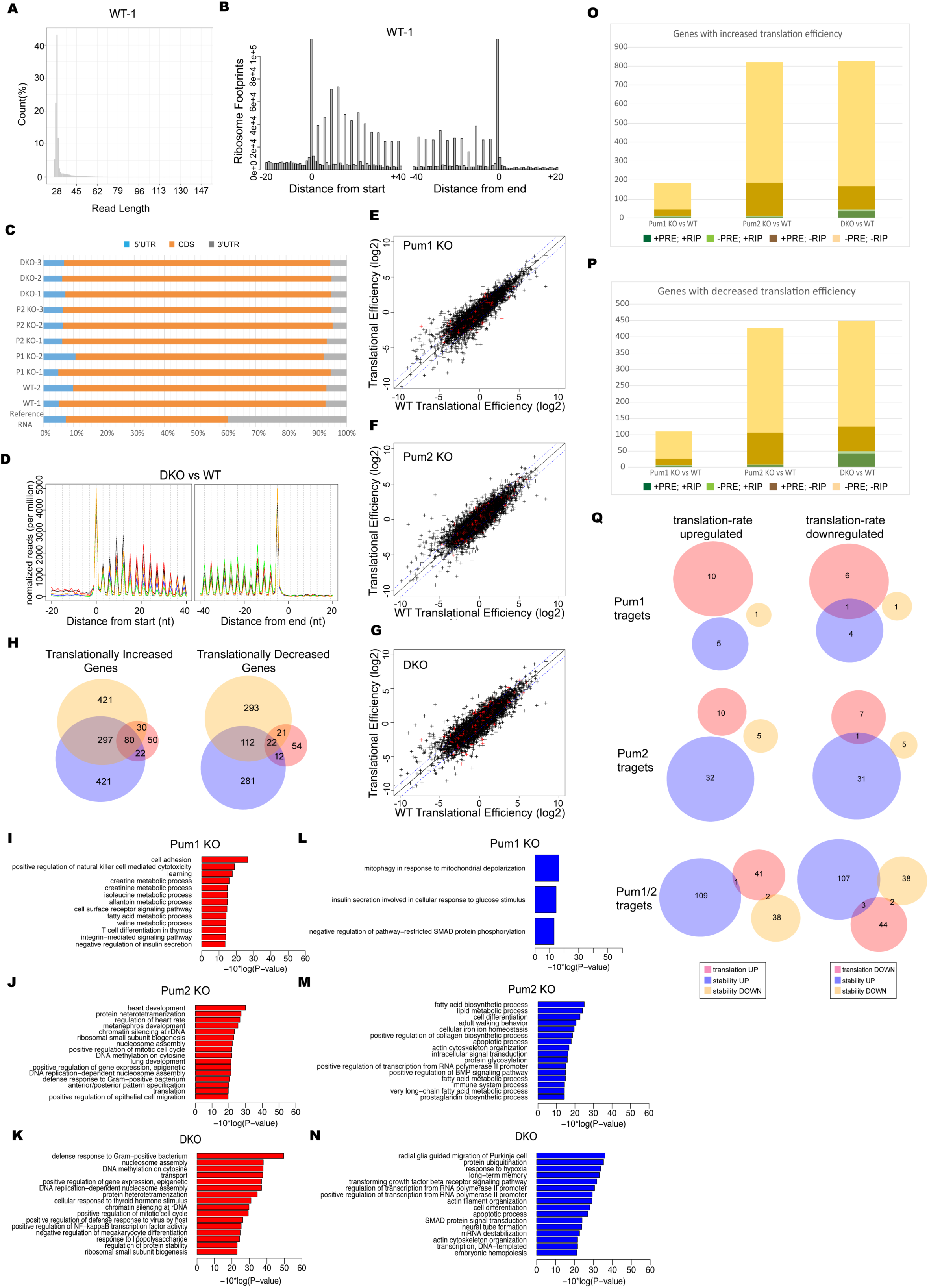
Pum1 and Pum2 regulate the translation of many mRNAs in ESCs. **(A)** Size profile of ribosomal footprints in wildtype ESCs from replicate one. y axis is the percentage, x axis is the length. All other samples from wildtype, *Pum1^−/−^*, *Pum2^−/−^*, and double mutant ESCs are extremely similar. **(B)** The metaplot of ribosomal P-site mapping around the start and stop codons of annotated protein coding regions in wildtype ESCs (replicate 1). The trinucleotide periodicity feature of ribosomal protection assay is clearly evident. +20 bp, -40bp region relative to start codon is displayed for translation starting region; +40bp, -20bp region relative to end codon is displayed for translation stopping region. **(C)** The percentage distribution of P-sites in 5’UTRs, CDS, and 3’UTRs in all genotypes.. **(D)** The metaplot of P-site mapping abound the start and the stop codon of annotated CDs in wildtype black curve) and *Pum* double mutant ESCs. Black and red curves correspond to wildtype samples; blue, green and orange curves correspond to *Pum* double mutant samples. Reads were normalized to sequencing depth. (**E-G**) Scatterplot of the translation efficiency of mRNA isoforms in *Pum1*^−/−^ (E), *Pum2*^−/−^ (F), and double mutant (G) ESCs compared to that in wildtype. Only mRNA isoforms with FPKM>1 are included. Red colored mRNAs are Pum1-(E), Pum2-(F), and Pum1-and/or-Pum2 (G) target RNAs by RIP-ChIP. The diagonal black line is y=x line; blue dashed lines are y=x+1 and y=x-1 lines. (H) Venn diagrams of translationally increased (left diagram) and decreased (right diagram) genes in *Pum1*^−/−^, *Pum2*^−/−^ and *Pum* double mutants against wildtype. (**I-K**) Top sixteen GO terms enriched in translationally-increased mRNA genes in *Pum1*^−/−^ (I), *Pum2*^−/−^ (J) and double mutant (K) ESCs, according to the *p* value. (**L-N**) Top sixteen GO terms enriched in translationally decreased mRNA genes in *Pum1*^−/−^ (L), *Pum2*^−/−^ (M) and double mutant (N) ESCs, according to the *p* value. (**O, P**) The statistics of direct target and PRE-containing mRNA isoforms that are in translationally increased (O) or decreased (P) in *Pum1*^−/−^, *Pum2*^−/−^ and *Pum* double mutants against wildtype. (Q) Venn diagram illustrating the overlap among translation-regulated Pum target genes and stability-regulated Pum target genes.

We next analyzed changes in mRNA translation in *Pum1^−/−^*, *Pum2^−/−^* and the double mutant ESCs. The ribosome distribution pattern in mutant ESCs was similar to that of wild type cells; ribosomal stalling was not observed in wild type or mutant ESCs (Figure 7D, Supplemental Figure 11 J, K), indicating that Pum1- and/or Pum2-deficiencies do not affect the dynamics of translation. In *Pum1^−/−^* and *Pum2^−/−^* ESCs, 182 and 820 genes, respectively, show increased translation; whereas 109 and 427 genes, respectively, show decreased translation (Figure 7E, F). In the double mutant ESCs, 828 genes display increased translation; whereas 448 genes show decreased translation (Figure 7G). The up- and down-regulated mRNAs in the three types of mutant ESCs partially overlap (Figure 7H), consistent with the overlapping function of Pum1 and Pum2. Gene ontology analysis revealed that the up-regulated genes in mutant ESCs are enriched with nucleosome assembly, DNA methylation and epigenetic regulation (Figure 7I-K). The down-regulated genes in mutant ESCs are enriched in ubiquitination, response to hypoxia, long-term memory, and transcriptional pathways (Figure 7L-N). These findings reflect the diversity of genes regulated by Pum1 and Pum2.

### Pum proteins repress or activate translation and turnover of different subsets of target mRNAs

To investigate how Pum proteins regulate the translation of their direct target genes, we examined the change in translational efficiency of Pum1– and/or Pum2-bound mRNAs in their corresponding mutant and double mutant ESCs. In *Pum1^−/−^* and *Pum2^−/−^* ESCs, 10 (0.7%) and 10 (2.6%) of 1461 Pum1 and 379 Pum2 targets, respectively, are translationally up-regulated (Figure 7O). In *Pum1^−/−^*; *Pum2^−/−^* ESCs, only 44 (3%) of the 1486 direct Pum1- and/or Pum2-bound mRNAs are translationally up-regulated (Figure 7O). These results indicate that Pum1 and Pum2 only suppress small subsets of their target mRNAs, in contrast to the common belief that Pum proteins regulate all of their target genes.

Similarly, only 7 (0.5%) and 8 (2.1%) of 1461 Pum1 and 379 Pum2 target mRNAs are translationally down-regulated in *Pum1^−/−^* and *Pum2^−/−^* ESCs, respectively (Figure 7P). In *Pum1^−/−^*; *Pum2^−/−^* ESCs, only 49 (3.3%) of 1486 Pum1- and/or Pum2-bound mRNAs are translationally down-regulated (Figure 7P). These results indicate that Pum1 and Pum2 can promote the translation of a similar number of their target mRNAs, in contrast to the well-known function of Pum proteins in translational repression.

We then investigated whether Pum1 and Pum2 regulate the stability of their direct targets. In *Pum1^−/−^* and *Pum2^−/−^* ESCs, 1 (0.07%) and 5 (1.32%) of 1461 Pum1 and 379 Pum2 targets, respectively, become less stable; whereas in *Pum1^−/−^*; *Pum2^−/−^* ESCs, only 40 (2.70%) of the 1486 direct Pum1- and/or Pum2-bound mRNAs become less stable. In *Pum1^−/−^* and *Pum2^−/−^* ESCs, 5 (0.34%) and 32 (8.44%) of 1461 Pum1 and 379 Pum2 targets, respectively, become more stable; whereas in *Pum1^−/−^*; *Pum2^−/−^* ESCs, only 110 (7.40%) of the 1486 direct Pum1- and/or Pum2-bound mRNAs become more stable. There is little overlapping between translationally regulated and stability-regulated target mRNAs (Figure 7Q), indicating that Pum proteins exerts a single mode of regulation towards an individual target mRNA.

## DISCUSSION

### Pum1 and Pum2 together are essential for early embryogenesis by initially regulating ESC pluripotency at the morula stage

Pum1 mRNA is expressed more ubiquitously in fetal and adult tissues than Pum2 mRNA (28). Consistent with this, we observed that Pum1-deficient mice have a more severe phenotype than Pum2-deficient mice. *Pum1*^−/−^ mice display significantly decreased body weight and have uniformly smaller organs, but *Pum2*^−/−^ mice only exhibit slightly lower body weights (1.7 – 4.4g less) than wild type littermates, validating previous reports (22) (26). Thus, Pum1 plays a more important role than Pum2 in embryogenesis. The partial redundancy between Pum1 and Pum2 indicate overlapping functions, which together are essential for embryogenesis, since knocking out both Pum proteins causes developmental delay starting at e3.5, when ESCs are established, and lethality by e8.5. More importantly, our molecular analysis of the pluripotency and the development of three germ layers of the double mutant embryos reveal that the embryonic lethality may initially be due to ESC defects at e3.5 in self-renewal and differentiation—two complementary aspects of pluripotency--followed by abnormal differentiation and hypoplasia during germ layer formation, ultimately resulting in a disorganized and inviable embryo by ∼e7.5-8.5.

### Pum1 and Pum2 have seemingly opposing but complementary functions in ESC pluripotency

In support of the above conclusion, Pum1 and Pum2 have important functions in ESCs. Remarkably, despite their extreme high homology and overlapping targets, Pum1 and Pum2 appear to have very different functions in ESC pluripotency, with Pum1 promoting differentiation and Pum2 promoting self-renewal. The differentiation–promoting function of Pum1 is particularly intriguing since previous studies in invertebrate models have only implicated Pum proteins in stem cell-self-renewal. Its role in differentiation is, however, consistent with the study by Leed, *et al*. (31), in which haploid ESCs generated an increased number of AP-positive (undifferentiated) colonies after Pum1 knockdown. In our study, Pum1 deletion in diploid ESCs not only increased the number of AP-positive colonies but also the total colony number, with the percentage of undifferentiated colonies remaining similar to that of wild type ESCs. Therefore, Pum1 promotes not only the differentiation of ESCs, but also inhibit ESC proliferation. This function is also opposite to known function of Pum proteins in previous studies in diverse model systems.

In contrast to Pum1-deficient ESCs, Pum2-deficient ESCs fail to maintain the expression of pluripotency genes but instead precociously express differentiation genes of the three germ layers after being transferred to feeder- and/or LIF-free condition. However, such ESC defects did not lead to correspondingly detectable defects in embryogenesis. This could be due to the much more stringent *ex vivo* condition that promotes differentiation, as compared to the supportive *in vivo* environment, since it is known that a mutation can generate strong ESC phenotype but not *in vivo* (36). In any case, the complementary functions of Pum1 and Pum2 must be due to the unique targets of their own that generate unique combinatorial effect of regulation among themselves and with other common targets.

### Pum1 and Pum2 negatively regulate each other and might auto-regulate its own expression

Pum1 and Pum2 bind to each other’s mRNA to suppress their translation. Thus, the effect of knocking out one Pum might be partially compensated by the over-expression of the other, possibly through enhanced regulation of their common targets and/or ectopic binding to mRNAs that are normally only bound by the other Pum. This strategy should have a survival advantage. In addition, this negative inter-regulatory feedback loop may fine-tune the steady-state levels of Pum1 and Pum2 in the cell, which in turn precisely controls the level of expression of its target genes such as pluripotency genes. Such fine-tuning is crucial for ESC function and embryogenesis.

Furthermore, we found that Pum1 and Pum 2 bind to their own mRNAs. These observations, consistent with previous reports that human PUM2 protein binds to Pum2 mRNA (37), indicate that PUM proteins can auto-regulate their own expression. Indeed, *Drosophila* Pum participates in a negative-feedback mechanism with Nanos to protect neurons from overactivity of Pum (38). Hence, we propose that Pum1 and Pum2 negatively regulate their own expression.

### Pum1 and Pum2 together are master regulators of embryogenesis

PUF proteins are described as “regulators of regulators” (39) because they control the expression of transcription factors and kinases that have many diverse regulatory effects on gene expression. This study supports this notion. The identification of 1461 target mRNAs of Pum1 and 379 target mRNAs of Pum2 in mouse ESCs is consistent with the number of known Pum targets in human HeLa cells (40) and in the mouse testis (25). It is interesting to note that among mRNAs bound by both Pum1 and Pum2, many of them encode functionally related proteins. These data are not only consistent with previous reports of overlapping sets of mRNA bound to human PUM1 and PUM2 (33), but also reveal that Pum2 binds to an almost complete subset of Pum1 mRNA targets. The 354 common mRNA targets of Pum1 and Pum2 are particularly intriguing, since these may represent the more evolutionarily conserved targets of Pum proteins.

The analysis of all Pum target mRNAs reveals that Pum1 and Pum2 are indeed global regulators of gene expression in ESCs and early embryos, as evident by the finding that transcription factors are among the most highly enriched categories of target mRNA in these cells. This also likely explains why Pum-target mRNAs are only a small subset of mRNAs that are affected in Pum-deficient ESCs. It is conceivable that Pum proteins represses or activate transcription factors, which then regulate many other genes. Other highly enriched Pum targets encode proteins involved in embryonic patterning. Reduced self-renewal capacity of *Pum1*^−/−^; *Pum2*^−/−^ ESCs and early embryonic lethality of the *Pum1*^−/−^; *Pum2*^−/−^ mice indicate that the dysregulation of these targets have significant consequences. The delicate balance between self-renewal and differentiation, and of gene activation versus repression must occur for embryogenesis. This balance is incredibly complex, and Pum proteins are “master regulators” that coordinately regulate diverse cellular processes through the regulation of large number of target genes.

### Pum1 and Pum2 not only repress translation but also promote translation and repress or enhance the stability of different target mRNAs

A surprising finding from our study is that Pum1 and Pum2 repress the translation of only a few of their direct target mRNAs, respectively, in contrast to the expected repression towards most, if not all, target mRNAs. Even more remarkably, Pum1 and Pum2 promote the translation of similar percentages of the target mRNAs. Moreover, Pum1 and Pum2 can even stabilize or destabilize yet other sizable subsets of their direct target mRNAs. These novel and substantial modes of Pum regulation add to the already complex network of regulation and provide rich stream of future investigations to examine how Pum proteins have such different regulatory impact on different target mRNAs. Such studies should reveal novel molecular mechanisms of Pum-mediated gene regulation and how these mechanisms contribute to development.

## MATERIALS AND METHODS

### Mouse ESC and Blastocyst Culture

CCE cells (Stem Cell Technologies) were cultured on gelatin-coated tissue culture plates in the presence of sterile-filtered ESC media with leukemia inhibitory factor (LIF) (ESGro Millipore), ES-qualified FBS (Gemini Bioproducts), NEAA, L-Glut, Pen/Strep, and Sodium Pyruvate. Media was changed every day and the cells were passaged every other day with TrypLE Dissociation Reagent (Gibco). Embryoid bodies (EB) were generated by culturing ESCs on non-adherent tissue culture plates in the absence of LIF; embryoid bodies were collected every 48 hours for qRT-PCR analysis. Overnight blastocyst culture was in KSOM Embryo Media (Millipore).

### Isolation and analysis of *Pum* double mutant embryos

*Pum1flox; pum2flox; rtTA*[*ROSA*]*26-Cre* female mice were superovulated and individually mated with a male *pum1^flox/flox^; pum2^flox/flox^; rtTA*[*ROSA*]*26-Cre* mice. Detection of a copulation plug the following morning was used to confirm successful mating. 200mg/ml Doxycycline (D9891, Sigma) in 2% sucrose water was then fed to mice to produce *pum1^−/−^; pum2^−/−^* mice. Embryos at appropriate age were dissected for the following immunofluorescent staining. To obtain *Pum1^+/−^; Pum2^+/−^* control embryos, we crossed *C57BL/6J* female mice with *pum1^flox/flox^, pum2^flox/flox^; Dox-Cre* male mice, followed by the same Dox treatment. The genotypes were confirmed by immunofluorescence staining of Pum1 and Pum2 antibodies (Supplemental Figure 4C-F)

Embryos were fixed in 4% paraformaldehyde in PBS for 15 minutes, and then permeabilized in PBS 0.5% Triton X-100 for 10 minutes. After blocking with 5% NGS and 0.15% Tween 20 in PBS, embryos were stained with primary antibodies: anti-Oct4 (sc-5279, Santa Cruz), anti-Gata4 (sc-25310, Santa Cruz), anti-Cdx2 (sc-56818, Santa-Cruz), anti-Pum1 (AB92545, Abcam) and anti-Pum2 (AB92390, Abcam). Antibodies were diluted 1:200 in blocking buffer and embryos were incubated in the primary antibody solution at 4°C overnight, followed by the appropriate secondary antibodies. Secondary antibodies labeled with Alexa fluorophores (Invitrogen) were diluted 1:500 in blocking buffer. Nuclei were stained with DAPI (Molecular Probes). Embryos were mounted on glass slides with Vectashield (H-1000, Vector Labs). Images were captured with a confocal microscope (Leica SP5) and Leica confocal software. Embryos were genotyped by anti-Pum1 and anti-Pum2 antibody staining (Supplementary Figure 3).

### ESC Derivation and Embryoid Body Formation

Mice matings were timed by checking for plugs (Day 0.5). Pregnant females were sacrificed at e3.5, and blastocysts were flushed from uteri in warm M2 media. Blastocysts were incubated in KSOM overnight under mineral oil in standard cell incubator conditions (5% CO2), then plated on MEFs the following day for passage. The blastocysts were allowed to attach to the plates for 3 days, then the media was changed daily for another 3 days until the cells were passaged between day 7-10. All newly derived cell lines were assayed for Nanog and Oct 4 expression and were frozen at low passage number for use in all experiments. *Pum1^−/−^*,*Pum2^−/−^*, and their double mutant embryoid bodies were generated by removal of LIF and culturing in hanging drops for two days, then on non-adherent tissue culture plates. Cells were collected every 48 hours up to 14 days for quantitative RT-PCR for markers of pluripotency as well as endodermal, mesodermal, and ectodermal lineages.

### Antibodies

The following antibodies were used: Pum 1 polyclonal goat anti-mouse antibody (Novus), Pumilio 2 polyclonal rabbit anti-mouse antibody (Novus), Pum 1 monoclonal rabbit anti-mouse antibody (Epitomics), Pum 2 rabbit anti-mouse monoclonal antibody (Epitomics), Cyclin B1 polyclonal rabbit anti-mouse (Cell Signaling), Cyclin E2 polyclonal rabbit anti-mouse (Cell Signaling), Suz 12 polyclonal rabbit anti-mouse (Cell Signaling), Cyclin A polyclonal mouse anti-human (Cell Signaling), CDK4 monoclonal mouse anti-human (Cell Signaling), Cyclin D1 monoclonal mouse anti-human (Cell Signaling), Sox2 goat anti-mouse (Santa Cruz), Nanog polyclonal rat anti-mouse (eBioscience), APC1 polyclonal rabbit anti-mouse (Bethyl), RENT1 polyclonal rabbit anti-mouse (Bethyl), ZNF198 polyclonal rabbit anti-mouse (Bethyl), RNA Pol II mouse anti-human (Abmart), GAPDH polyclonal rabbit anti-mouse (Sigma).

### Immunostaining

Mouse ESCs were grown on gelatin-coated tissue-culture treated plates, transferred to slides and fixed for 15 minutes in 4% paraformaldehyde in PBS. Immunofluorescence staining was performed with standard protocols using an Alexa Fluor 488 conjugated rabbit anti-mouse secondary antibody. A 1:1000 dilution of DAPI was incubated with fixed cells after staining to label the nuclei. Images were taken with a Leica TCS SP5 Spectral Confocal Microscope in the sequential scanning mode.

### Knockout mouse generation, mouse husbandry, and genotyping

Conditional *Pum1* and *Pum2* knockout mice were generated at the University of Connecticut Gene Targeting and Transfer Facility in Farmington, CT. *Pum1^Flox/+^* and *Pum2^Flox/+^* mice were mated with *EIIa-Cre* mice to drive the expression of Cre recombinase in pre-implantation embryos and generate knockout alleles. The *Pum1* knockout allele produces a truncated Pum1 protein of 448 amino acids that is upstream of the Pum-HD with the last 31 amino acids out-of-frame; the Pum2 knockout allele produces a 40 amino acid protein with the last 23 amino acids out-of-frame. Mice in which the floxed *Pum1* or *Pum2* allele was excised were confirmed by genotyping, and all subsequent crosses were performed on global *Pum1/2* heterozygous mice in the absence of Cre. For genotyping, tails were clipped at 21 dpp and digested overnight at 55 degrees with Proteinase K. Genotyping samples were diluted 1:10 with distilled water and 2 uL of each sample was used as template for PCR. The following genotyping primers were used for genotyping Pum1 and Pum2. Pum1: Lox gtF: atcaggttgccagtttcacc, Lox gtR: ttttcactgaaccagcaagg, Frt gtR: tgattctgcaaggacagcac, Pum2: Lox gtF: CATACTGTCACTAACCTGTC Lox gtR: TGCTGATCTCATCTTCGGAG, Frt gtR: ATGCATATGTGCCATGGAGC. Cre internal control primers: Fwd: CTA GGC CAC AGA ATT GAA AGA TCT, Rev: GTA GGT GGA AAT TCT AGC ATC ATC C, Cre primers Fwd: GCG GTC TGG CAG TAA AAA CTA TC, Rev: GTG AAA CAG CAT TGC TGT CAC TT.

### Nested PCR for genotyping blastocysts

Two rounds of PCR were performed for genotyping blastocysts. DNA was isolated from blastocysts using the QIAamp Investigator Kit (Qiagen), eluting in 23 uL of nuclease-free water. 10 uL of this DNA was used for each PCR. The following primer pairs were used for Round 1 and Round 2 reactions for Pum1 and Pum2. WT PUM1 Round 1: For - TGAACTTGGCTGTTGATGGGTCCG, Rev-TCCATAGACTGGTCCTCTGTCTCACCT. KO Pum1 Round 1: For - tgaacttggctgttgatgggtccg, Rev – agggtgtggccttttggccttt. WT Pum2 Round 1: For – ggtttccttgctctttagtcttcccca, Rev – cggtcgttactctctatccctgacac. KO Pum2 Round 1: For - ggtttccttgctctttagtcttcccca, Rev – accatgctccaaagtcccacgg. WT Pum1 Round 2: For – Tgaacaggtgaattgctgactgtgaa, Rev – Cctggctttgggagtttcagggc. Pum1 KO Round 2: For – Tgaacaggtgaattgctgactgtgaa, Rev – Agggtgtggccttttggcct. WT Pum2 Round 2: For – tgcttttgttcatggttttctg, Rev – gcatgctgatctcatcttcg. Pum2 KO Round 2: For – tgcttttgttcatggttttctg, Rev – ctgcttccacttttgcatga.

### Transfection and siRNA knockdown

Cells were transfected with Lipofectamine 2000 at a ratio of 1.5 uL Lipofectamine to 5 uG DNA to 45 uL OptiMEM (Invitrogen). Reverse transfections were performed in order to maximize transfection efficiency, and repeat transfections were performed on Day 2 and 4 for Cre-mediated excision of floxed alleles. siRNA transfections were performed according to the same protocol, and were repeated 48 hours after the first transfection for increased knockdown efficiency. Pum1 siRNA (Dharmacon Smart POOL) contained a 5nmole mixture of 4 different siRNA sequences. Non-silencing siRNA (scrambled) was used as a control for all siRNA experiments. Luciferase reporter construct transfections were completed as above but using 1- 10ng Firefly luciferase plasmid (experimental) and 100pG Renilla luciferase (control).

### RNA-immunoprecipitation-microarray (RIP-Chip)

Mouse ESCs were lysed in cold MLB buffer (150mM NaCl, 0.1% NP-40, 0.1% Triton-X, ph 7.4) with 1X complete mini EDTA-free protease inhibitor cocktail (Roche). Total cellular lysate was spun at 10,000 RPM for 15 minutes, and supernatants were used for all subsequent experiments. The total lysate was pre-cleared with equilibrated Protein A Sepharose beads (GE Healthcare) for 2 hours at 4 degrees. Pre-cleared lysate was added to primary antibody for 4-6 hours at 4 degrees, and then Protein A beads were added and nutated at 4 degrees for an additional 2 hours. RNA was isolated directly from the beads using Trizol, and was further purified using a QIAquick kit (Qiagen). Samples were amplified and hybridized to an Illumina mouse Ref v2 BeadChip array (Supplemental Figure 9A, B). Experiments were conducted in triplicate, and a negative control using blocking peptide against which the antibody was raised was used for the normalization of all data. The immunoprecipitated mRNAs were normalized to control mRNAs using loess regression and quantile normalization, and the p value and fold enrichment of Pumilio mRNA targets were calculated, plotted, and visualized by heat map and volcano plots. Gene Set Enrichment Analysis (GSEA) was conducted for mRNA targets of Pumilio to determine significant functional enrichments of particular pathways. Gene Ontology annotations (GO Term), modified Fisher Exact p-values, and the DAVID Clustering procedure were utilized to measure the similarity of all gene-gene pairs.

### Statistical analysis

Error bars represent means ± SE; differences between means were calculated using the Student’s t-test and/or analysis of variance. p values < 0.05 were considered statistically significant (as marked by asterisks).

### Bioinformatic analysis of the transcriptomes and ribosomal protection assays of wild-type and mutant ESCs

#### 1. Evaluation of sequencing quality. FastQC

(https://www.bioinformatics.babraham.ac.uk/projects/fastqc/) was implemented to evaluate the quality of raw sequence data from Illumina high throughput sequencing machine.

#### 2. Sequence trimming and mapping

Ribo-seq and mRNA-seq data were trimmed off the 3’ end linker sequence by cutadapt1.16 (DOI:10.14806/ej.17.1.200) with -e 0, -m 27 parameters to allow for zero error rate and only sequences longer than 26nt were kept. We also kept the sequences if the linker could not be found. mRNA-seq sequences were trimmed under paired-end mode, while Ribo-seq sequences under sing-end mode (R1 reads were used). We aligned the trimmed and size-selected Ribo-seq data (R1 reads) to mm10 genome along with the GRCm38.81 gene annotation, using alignment software STAR2.7. Up to 2 mismatches were allowed and sequences with multiple mappable locations were excluded. Length distribution of mappable sequences from Ribo-seq was examined. Mapped reads of 27-29 nucleotides dominated in the libraries (Figure 7A).

For the mRNA-seq data, we aligned the trimmed and size-selected mRNA-seq data to mm10 genome along with the GRCm38.81 gene annotation using alignment software STAR2.7 with paired-end mode. Up to 2 mismatches were allowed and sequences with multiple mappable locations were excluded. Alignment bam files were collected for downstream analysis.

#### 3. Transcriptome(mRNA-seq) data analysis

We used Cufflinks (version 2.2.1) (Trapnell et al., 2012) to estimate the abundance of each transcript and to test for differential expression across RNA-Seq samples. Upper-quartile-norm was applied for normalization to improve the robustness of differential expression calls for less abundant genes and transcripts. Ensembl gene annotation GRCm38.81 was used for calculating gene expression level. The correlation between pairs of replicates for each condition was evaluated by Spearman correlation based on gene expression level (Supplementary Figure 11L). The mRNA isoforms/genes with fold change >=1.5 and p-value<=0.05 against control conditions were defined as differentially expression isoforms/genes and displayed in Heatmaps (Supplemental Figure 10A-F).

#### 4. Ribosomal profiling data analysis

The correlation between replicates for each condition was evaluated by Spearman correlation based on gene expression level (Supplementary Figure 10L). We applied riboWaltz (41), an R package to calculate the P-site offset and perform the diagnostic analysis/inspection of the ribosomal profiling data. We examined the percentage of P-sites falling in the three annotated transcript regions (5’ UTR, CDS and 3’ UTR), which revealed that the majority of the ribosome footprint reads were in CDS (Figure 7F).

The P-site offset was defined as the distance between the extremities of a read and the first nucleotide of the P-site itself. The majority of the Ribo-seq data were 27-35nt (89.9% in flox Pum1 rep1, 84.6% in flox Pum1 rep2, 98.0% in Pum1KO rep1, 85.6% in Pum1KO rep2, 89.6% in Pum2KO rep1, 98.8% in Pum2KO rep2, 97.4% in Pum2KO rep3, 95.3% in Pum1/2 double KO rep1, 95.3% in Pum1/2 double KO rep2, 95.5% in Pum1/2 double KO rep3). A representative size profile of flox Pum1 rep1 is shown in Figure 7A. Given this, we selected 27-35nt mappable reads for calculating the P-site offset, i.e. length_filter_mode = “custom” and length_filter_vector = 27:35.

To compare the P site profile between different conditions, we first normalized the signal against the sequencing depth. Then, the translational efficiency was calculated by RiboDiff (42) as

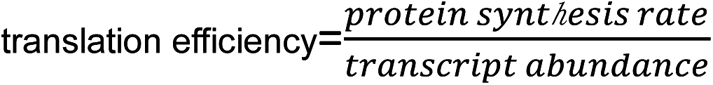

where the protein synthesis rate was defined as the normalized accumulated ribosomal signal in the CD region of transcript. The gene with translation efficiency fold-change >=1.5 and p-value<=0.05 was defined as an affected gene (Figure 7H-N).

#### 5. Gene Ontology enrichment (GO) analysis

GO analysis was performed using DAVID Bioinformatics resources 6.8 (43). P-value was calculated by Fisher’s exact test to measure the statistical significance of enrichment of the genes associated with GO terms.

## ACKNOWLEDGEMENTS

We would like to thank Dr. Natalia Ivanova for advice on the embryoid body differentiation assay, Dr. Apoorva Tewari for whole-mount Pum1 and Pum2 RNA in situ hybridization analysis of e8.5 and e9.5 embryos, Dr. Jing Xia for initial bioinformatics analysis, Min Young Lee for technical help with ESC derivation, Emily Mis for assistance with *in situ* hybridizations, Dr. Carmen Booth for sectioning and phenotypic analysis of mice, Dr. Shangqin Guo for the rtTA-Rosa26 mice, members of the Lin lab for helpful discussions and reagents, and Mr. Duy Phan for critical reading of the manuscript. K.U. was supported by NIH Medical Scientist Training Program Training Grant T32GM007205 and an American Fellowship from AAUW. This work was initially supported by the Mathers Award and a ShangahiTech University start-up fund and then by NIH R01 GM121386 (which did not overlap with the Mathers and ShanghaiTech support), to H.L.

