## Supplementary Information for "Pumilio Proteins Exert Distinct Biological Functions and Multiple Modes of Post-Transcriptional Regulation in Embryonic Stem Cell Pluripotency and Early Embryogenesis"

### SUPPLEMENTARY RESULTS

#### *Pum1*$^{-/-}$ mice are viable to adulthood but are smaller than their wild type littermates during fetal development

To further investigate how early the size difference between the Pum mutants and their wildtype siblings, we examined the morphology of e9.5 and e12.5 embryos from *Pum1*$^{-/-}$*; Pum2*$^{-/-}$ self-mating. *Pum1*$^{-/-}$*; Pum2*$^{+/-}$ embryos recovered at e9.5 showed developmental delay and posterior hypoplasia, with normal-appearing heart morphology and somite development (Supplemental Figure 3A). *Pum1*$^{-/-}$*; Pum2*$^{+/-}$ embryos were smaller than their littermates at e12.5, but showed no obvious developmental defects (Supplemental Figure 3B). Yolk sacs from e12.5 *Pum1*$^{-/-}$*; Pum2*$^{+/-}$ embryos were darker and more friable than yolk sacs from all other genotypes (Supplemental Figure 3B; lower right panel), suggesting that both defects in the supporting structures and embryonic tissue may be contributing to the phenotype of *Pum*-deficient embryos.

**PRE is important but not essential for translational repression or activation of Pum-target mRNAs**

Among the 10 translationally upregulated Pum1-bound genes in *Pum1*$^{-/-}$, 8 (80%) have one or more PRE motifs, while among 172 non-Pum1-bound but translationally up-regulated genes , 34 (20%) have at least one PRE motif (Figure 7O). Likewise, among 10 translationally upregulated Pum2-bound genes in *Pum2*$^{-/-}$, 8 (80%) have one or more PRE motif, while among 810 non-Pum2-bound but translationally up-regulated genes, 175 (22%) have at least one PRE motif (Figure 7O). Furthermore, among 43 translationally upregulated Pum1- and/or Pum2-bound genes in *Pum1*$^{-/-}$; *Pum2*$^{-/-}$, 35 (81%) have one or more PRE motifs, while among 785 non-Pum-bound but translationally up-regulated genes, only 125 (16%) have at least a PRE motif (Figure 7O). These results together indicate that PRE is important but not essential for translational repression of Pum-target mRNAs.

Notably, among the 7 translationally down-regulated Pum1-bound genes, 5 (71%) have one or more PRE motif, while among 102 non-Pum1-bound but translationally down-regulated genes, 19 (19%) have at least one PRE motif (Figure 7P). Likewise, among 8 translationally down-regulated Pum2-bound genes, 6 (75%) have one or more PRE motifs, while among 419 non-Pum2-bound but translationally down-regulated genes, 98 (23%) have at least one PRE motif (Figure 7P). Among 49 translationally down-regulated Pum1- and/or Pum2-bound genes, 41 (84%) have one or more PRE motifs, while among 399 non-Pum-bound but translationally down-regulated genes, 76 (19%) have at least a PRE motif (Figure 7P). These results together indicate that PRE might be important but not essential for translational activation of Pum-target mRNAs as well.

### SUPPLEMENTARY FIGURE LEGENDS

**Supplemental Figure 1 *Pum1* knockout and phenotype.**

**(A)** Conditional knockout strategy for Pum1. Exons 8 and 9 were flanked with loxP sites, and mice were mated with EIIa-Cre mice to generate global Pum1 knockout mice. **(B)** Genotyping of Pum1 +/- and Pum1 -/- mice. **(C)** Expected and observed ratios of Pum1 heterozygote matings. **(D)** Organ weight (in grams) of wild type (black bars), Pum1 +/- (dark gray bars), and Pum1 -/- (light gray bars) mice. **(E)** Normalized organ weight (in grams) compared to wild type mice (black bars normalized to 1).

**Supplemental Figure 2 *Pum2* knockout and phenotype.**

**(A)** Conditional knockout strategy for Pum2. Exon 3 was flanked with loxP sites, and mice were mated with EIIa-Cre mice to generate global Pum2 knockout mice. **(B)** Genotyping of Pum2 +/- and Pum2 -/- mice. **(C)** Expected and observed ratios of Pum2 heterozygote matings. **(D, E)** Body weight (in grams) of female (D)and male (E) of wild type (solid line), Pum2 +/- (dashed line), and Pum2 -/- (dotted line) littermates at 1, 7, 14, 21, and 28 days post partum (dpp), with bars indicating SEM. **(F)** Phenotype of wild type (wt), *Pum2*$^{+/-}$, and *Pum2*$^{-/-}$ littermates at 28 days post partum (dpp).

**Supplemental Figure 3 Embryonic phenotype of various combinations of *Pum1* and *Pum2* knockout.**

**(A)** Phenotype of e9.5 embryos with genotype indicated in lower left. Increased *Pum* deficiency results in proportionately smaller embryos; no *Pum1*$^{-/-}$; *Pum2*$^{-/-}$ double knockout embryos were recovered at this stage. **(B)** Phenotype of e12.5 embryos; genotype indicated in upper left. Pum-deficient embryos remain proportionately smaller at this stage, with *Pum1* $^{-/-}$; *Pum2*$^{+/-}$ yolk sacs appearing smaller and darker than wild type yolk sacs (lower right panel). No *Pum1*$^{-/-}$; *Pum2*$^{-/-}$ embryos were recovered at this stage.

**Supplemental Figure 4 Phenotype of e3.5-e4.5 *Pum1-Pum2* double mutant embryos.**

**(A)** e3.5 blastocysts flushed from uteri of *Pum1*$^{+/-}$;*Pum2*$^{+/-}$ x *Pum1*$^{+/-}$;*Pum2*$^{+/-}$ mated mice. two of the four *Pum1*$^{-/-}$; *Pum2*$^{-/-}$ blastocysts obtained appear morula-like (asterisks).

(**B**) Embryos, including the two embryos, were incubated overnight in KSOM Embryo Media (Millipore) at 37 degrees, and by e4.5 all appear to have formed blastocysts.

(**C-F**) Confocal images of e4.5 wildtype (WT, C, E) and the mutant embryos (D, F) immunostained for Oct4 (Gray), Gata4 (green), and Pum1 (red) or Pum2 (red). Nuclei were counterstained with DAPI (blue). Very low or no expression of Pum1 or Pum2 expression in the mutant embryos validates their genotype (C, E). Scale bars, 25µm.

**Supplemental Figure 5 Lineage differentiation in e3.5-e5.5 *Pum1-Pum2* double mutant embryos.**

(**A, B**) Development of trophectoderm in mouse embryos. Confocal images of WT (A) and *Pum1*$^{-/-}$;*Pum2*$^{-/-}$ mutant embryos (B) isolated at 3e3.5 immunostained for DNA (using DAPI), and Oct4 (green) and Cdx2 (red). The mutant embryo showed morula morphology and defect in trophectoderm lineage development, as indicated by the no or low expression of Cxd2. Scale bars, 25µm.

(**C-G**) Confocal images of wildtype (C, E) and mutant (D, F, G) embryos at e4.5 (C-D) and e5.5 9E-G). Embryos were immunostained using antibodies against Cdx2 (red), Gata4 (green), Oct4 (gray) and nuclei were counterstained with DAPI (blue). Gata4-labeled primitive endoderm lineage consistently shows differentiation, whereas Cdx2-labeled trophectoderm lineage shows a largely normal development. However, In severe e.5.5 mutant embryos (G), all cell lineages are disorganized. Scale bars, 50 µm.

**Supplemental Figure 6 *Pum* mutant embryos exhibit abnormal morphology defects in e6.5 and e7.5 *Pum1-Pum2* double mutant embryos.**

(**A, B**) Immunostaining of Gata4 and Cdx2 of e6.5 wildtype (WT, A) and *pum* mutant (B) embryos. Wildtype embryo shows a normal egg cylinder morphology with prominent epiblast epithelium (A). However, the mutant embryos do not contain epiblast and PrE is mislocalized. Scale bars, 50 µm. (**C, D**) Immunostaining of Pum1 counterstained with DAPI in e7.5 embryos. Comparing to WT embryo (C), epiblast in mutant embryo (D) is disorganized and does not show a clear structure. Scale bars, 100 µm.

**Supplemental Figure 7 Genotype of *Pum* mutant ESCs and their cell proliferation and apoptosis assays.**

(**A-D**) Genotyping. *Pum1*$^{Flox/+}$ and *Pum1*$^{Flox/Flox}$ ESCs were derived from e3.5 blastocysts and transfected with pBabe-Puro-Cre plasmid to generate *Pum1*$^{+/-}$ and *Pum1*$^{-/-}$ cell lines, respectively. Knockout of wild type *Pum1* was confirmed by genotyping (A), quantitative RT-PCR (B) and western blot analysis (C), indicating that a truncated protein is not detectably expressed. Dox-inducible Cre is integrated into the genome of *Pum1*$^{Flox/Flox}$*; Pum2*$^{Flox/Flox}$ ESCs as a stable cell line. The exogenous Pum1 cDNA was cloned into pTRE vector, which was then packaged as lentivirus particles and transfected into the *Pum1*$^{Flox/Flox}$*; Pum2*$^{Flox/Flox}$ ESC with a Dox-inducible Cre transgene. Adding Dox induced the deletion of endogenous *Pum1* and *Pum2* genes as well as the expression of *Pum1* from the cDNA to compensate for the lack of Pum proteins to ensure better survival of the ESCs. *Pum1*$^{-/-}$*; Pum2*$^{-/-}$ ESCs are acquired by this strategy, and two independent lines were sued for assays. The deletion of the Pum genes was confirmed by Western blotting (D).

(**E, F**) Cell growth curves of four ESC lines in feeders or feeder-free conditions. The cell number was shown as mean ± SD. T-test, *p<0.05, ** p<0.01, *** p<0.001.

(**G, H**) Cell doubling time of the four ESC lines grown with or without feeder cells.

(**I, J**) FACS results of wild type and *Pum1*$^{-/-}$*; Pum2*$^{-/-}$ ESCs. Cell are sorted by Annexin-V-FITC and PI.

**Supplemental Figure 8 Teratoma assays on the differentiation potential of *Pum* mutant ESCs.**

(**A**) Teratomas segregated from subcutaneous injection of nude mice after 18 days of growth.

(**B**) Growth curve of teratomas. (**C**) Weight of teratomas. (**D**) Differentiated tissues in paraffin sections of teratomas presented by HE staining. (**E**) Sox1 staining showing primitive neuron tubes from the ectoderm. (**F**) Foxa2 staining showing tubular epithelium from the endoderm.

**Supplemental Figure 9 RNA-Immunoprecipitation and Microarray analysis of Pum1 mRNA targets.**

(**A, B**) RNA-Immunoprecipitation-Microarray (RIP-Chip) experiments were conducted on ESCs to identify the mRNAs bound to Pum1. RNA was isolated from the beads of Pum1 Immuno-precipitates (IPs) and run on an Agilent gel analyzer before (A) and after (B) amplification in collaboration with the Yale Keck Microarray Facility. rRNA is likely present in the pre-amplified RNA (dark bands), and 100-200bp RNAs appear amplified (dark smear) in the post-amplified samples. **(C)** Scatter plot of the log intensity of control IPs (x axis) and Pum1 IPs (y axis) before (left) and after (right) normalization. **(D)** MA plot of the mRNA enrichment in control vs. IP before (left) and after (right) normalization. **(E)** Volcano plot of the log fold of enrichment of IP vs. control (x axis) and the paired t-test p value (y axis); the top 500 hits of the RIP-Chip by both p value and fold enrichment (diagonal line from the upper right corner to the lower left corner of the volcano plot) were used in further statistical analyses. **(F)** Normalized luciferase expression of reporter constructs containing control luciferase plasmid (Luc+), plasmid with the full-length 3'UTR of Cyclin B1 (Cyclin), or with site-directed mutagenesis of UGU → AGA within 3 putative NRE sites within the 3'UTR of Cyclin B1 (Mut1, Mut 2, Mut 3).

**Supplemental Figure 10 Pum1 and Pum2 regulate the expression of many mRNAs in ESCs.**

(**A-F**) Heatmaps of the expression of up- (A) or down- (B) regulated genes in *Pum1*$^{-/-}$ mutant ESCs; up- (C) or down- (D) regulated genes in *Pum2*$^{-/-}$ mutant ESCs; and up- (E) or down- (F) regulated genes in *Pum1*$^{-/-}$;*Pum2*$^{-/-}$ mutant ESCs. FPKM values in (A-F) are $\log_2$ transformed for the plot. (**G**) Top seven GO terms enriched in up-regulated genes in *Pum1*$^{-/-}$ mutant ESCs, according to the p-value. (**H**, **I**) Top sixteen

GO terms enriched in up- (H) or down- (I) regulated genes in *Pum2*$^{-/-}$ mutant ESCs, according to the p-value. (**J, K**) Top sixteen GO terms enriched in up- (J) or down- (K) regulated genes in in the double mutant ESCs, according to the p-value.

**Supplemental Figure 11 Ribosomal profiling signatures of mRNAs in ESCs.**

(**A-I**) The metaplots of P-site mapping around the start and the stop codon of annotated CDs in wildtype (A), *Pum1*$^{-/-}$ (B, C), *Pum2*$^{-/-}$ (**D-F**), and the double mutant (**G-I**) ESC replicate samples, visualizing the striking trinucleotide periodicity feature of ribosomal protection assay. +20 bp, -40bp region relative to start codon is displayed for translation starting region; +40bp, -20bp region relative to end codon is displayed for translation stopping region. (**J**) The metaplots of P-site mapping abound the start and the stop codon of annotated CDs in wildtype and *Pum1*$^{-/-}$ mutant ESCs. black and red curves correspond to wildtype; blue, green curves correspond to *Pum1*$^{-/-}$ mutant. (**K**) The metaplot of P-site mapping abound the start and the stop codon of annotated CDs in wildtype and *Pum2*$^{-/-}$ mutant ESCs. black and red curves correspond to wildtype; blue, green and orange curves correspond to *Pum2*$^{-/-}$ mutant. (**L**) Correlation among replicates of reads of ribosome-protected fragments.

### Supplemental Figure 1

**A**

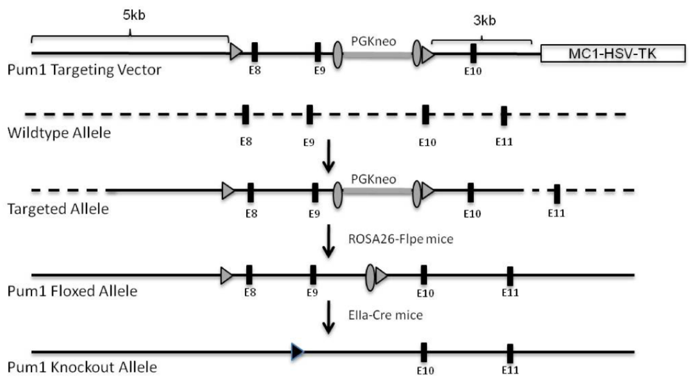


**B**

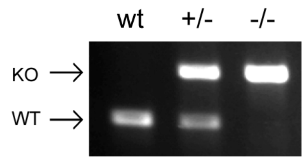


**C** *Pum1*$^{+/-}$ x *Pum1*$^{+/-}$

| | n | % |
|---|---|---|
| wild type | 68 | 28 |
| *Pum1* +/- | 142 | 58 |
| *Pum1* -/- | 34 | 13 |
| Total | 244 | |

**D** **E**

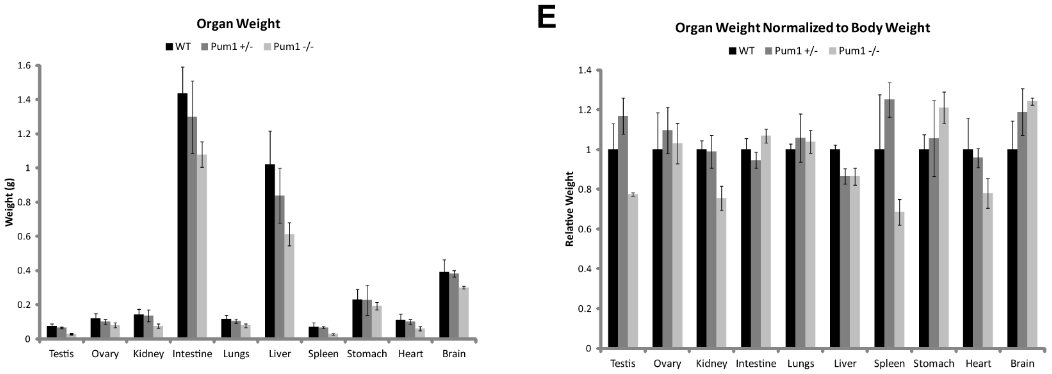

### Supplemental Figure 2

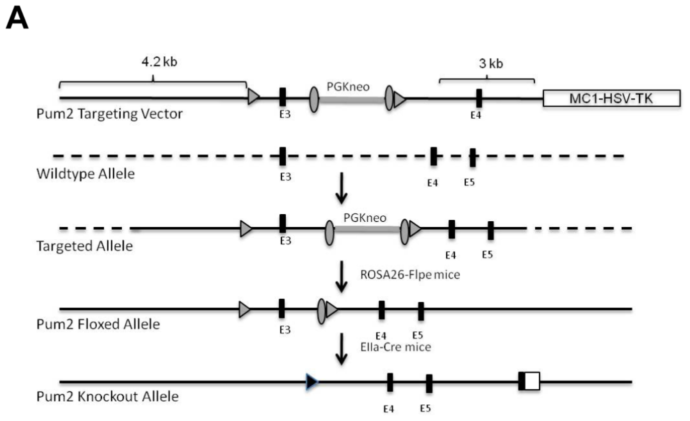


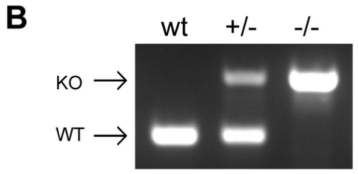


**C** *pum2*$^{+/-}$ x *pum2*$^{+/-}$

| | n | % |
|---|---|---|
| wild type | 42 | 26 |
| *pum2 +/-* | 76 | 47 |
| *pum2 -/-* | 43 | 27 |
| Total | 161 | |

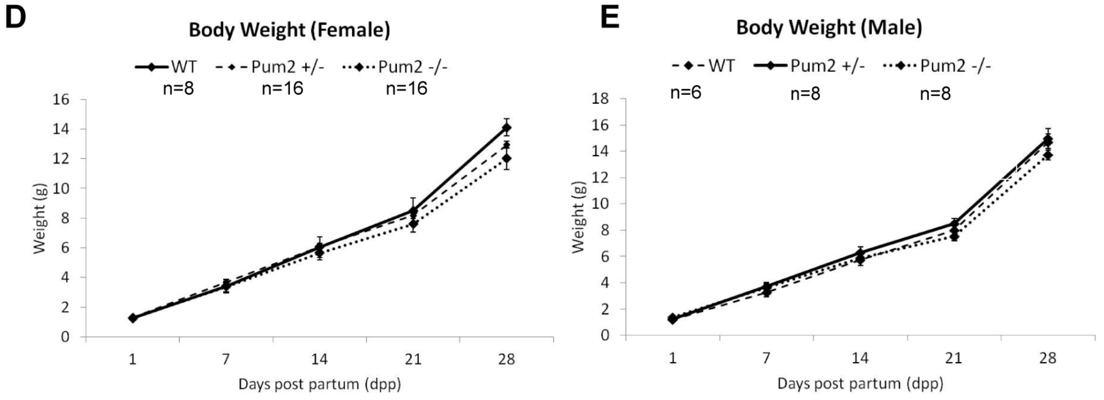


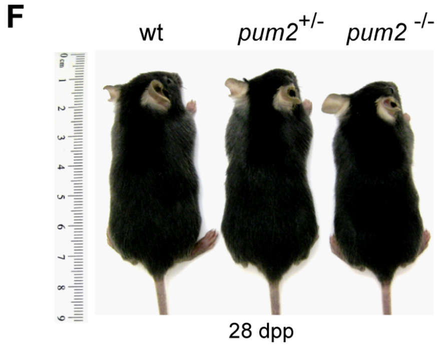

Supplemental Figure 3

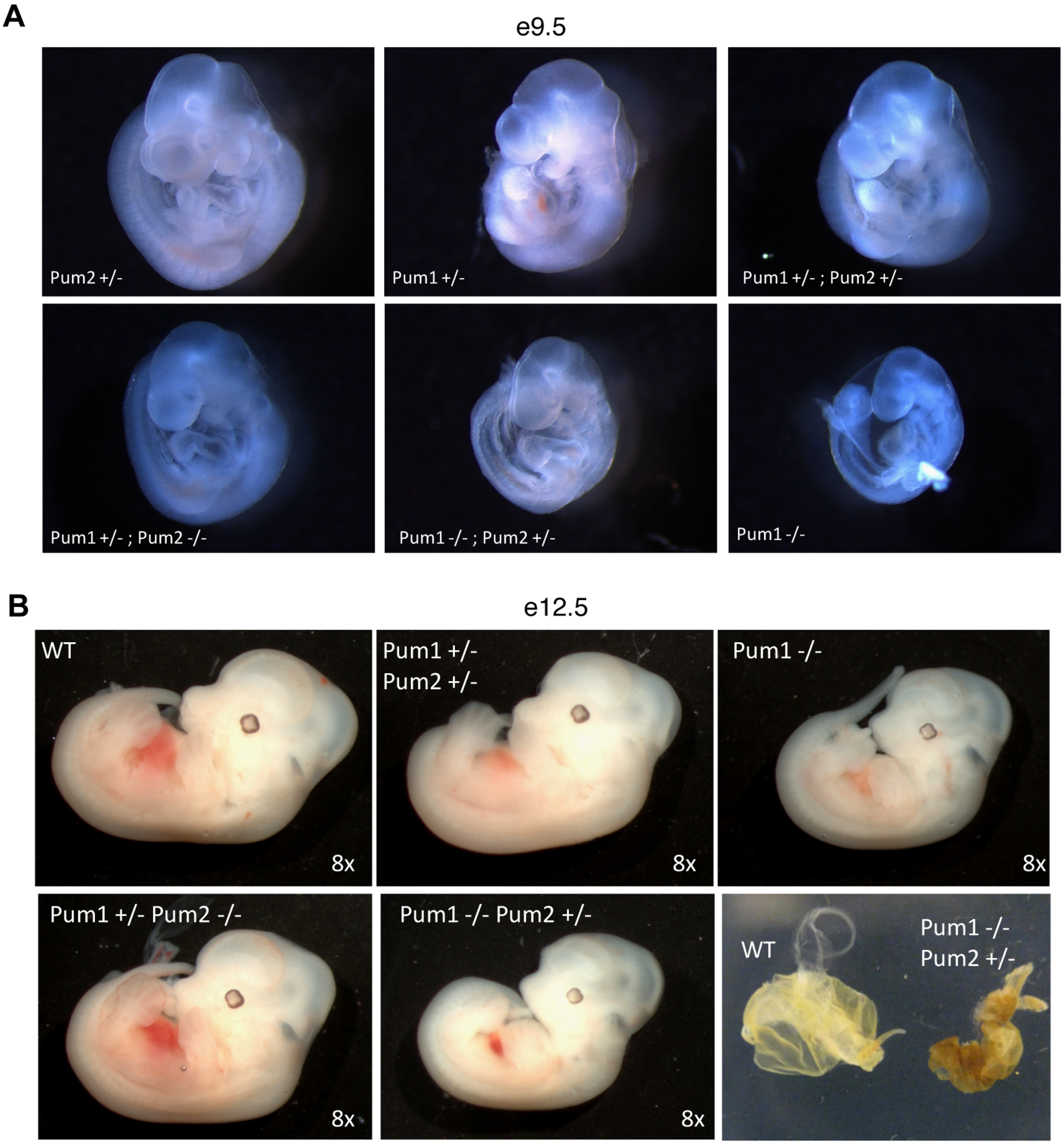

Supplemental Figure 4

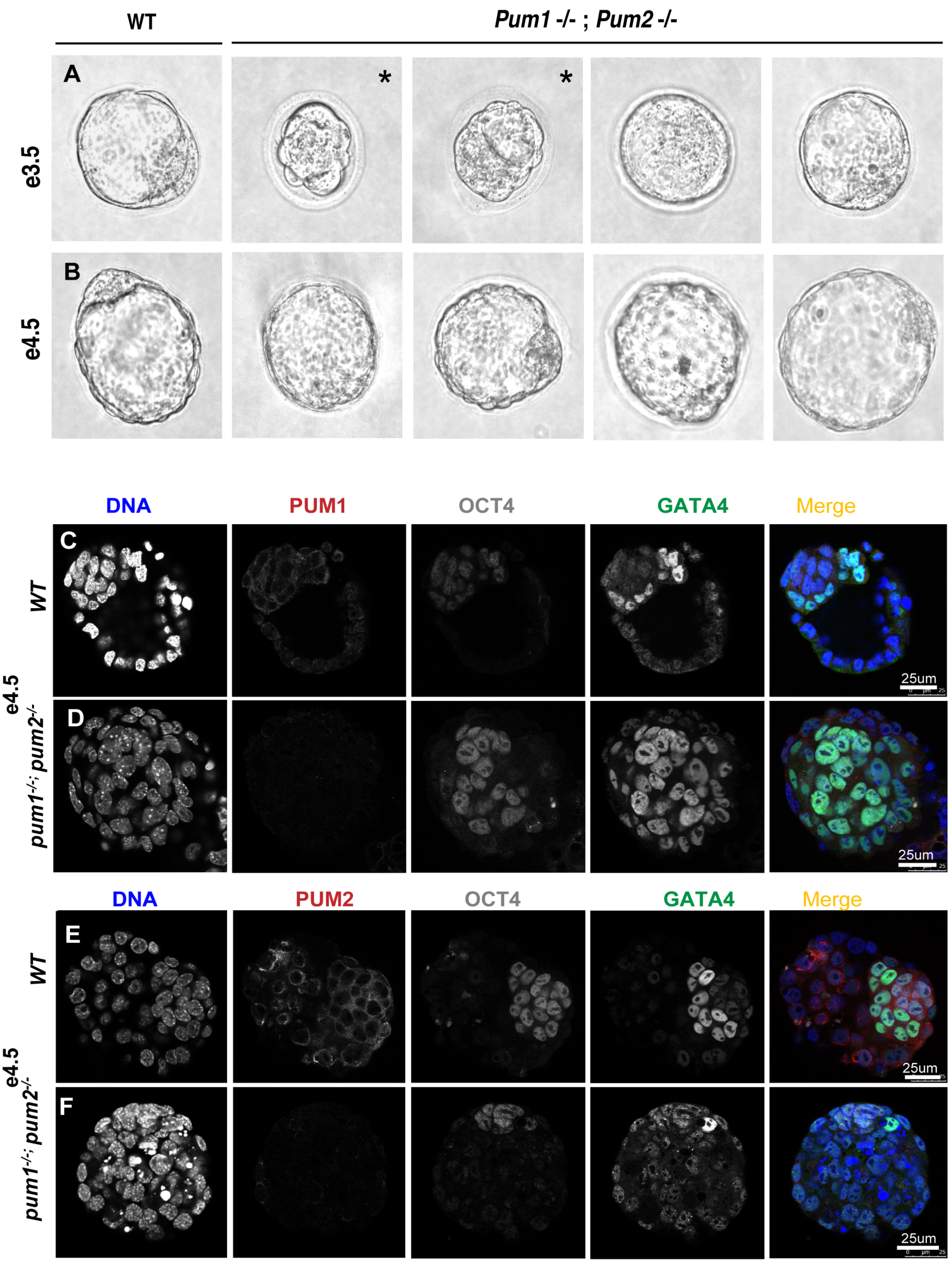

Supplemental Figure 5

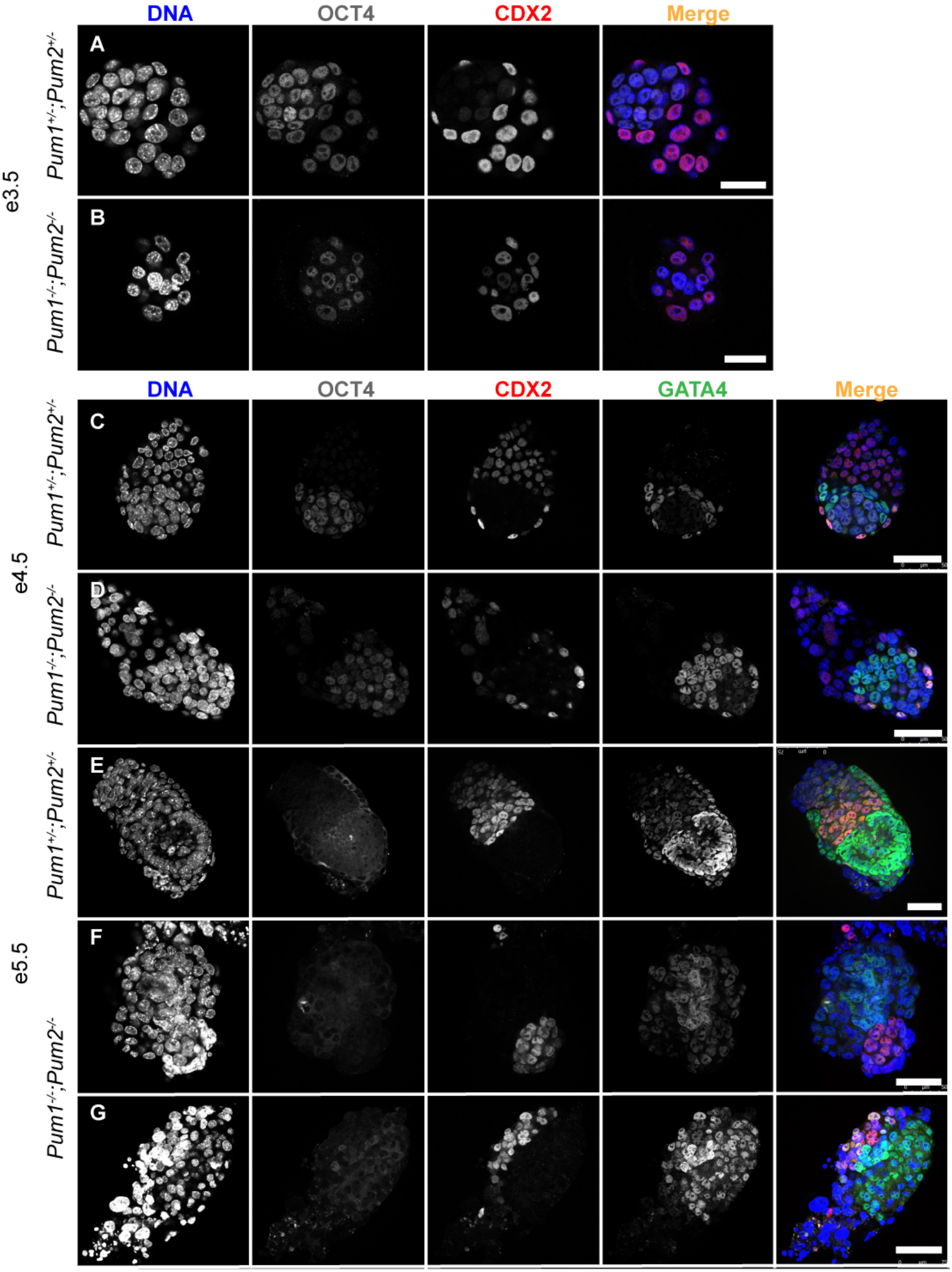

Supplemental Figure 6

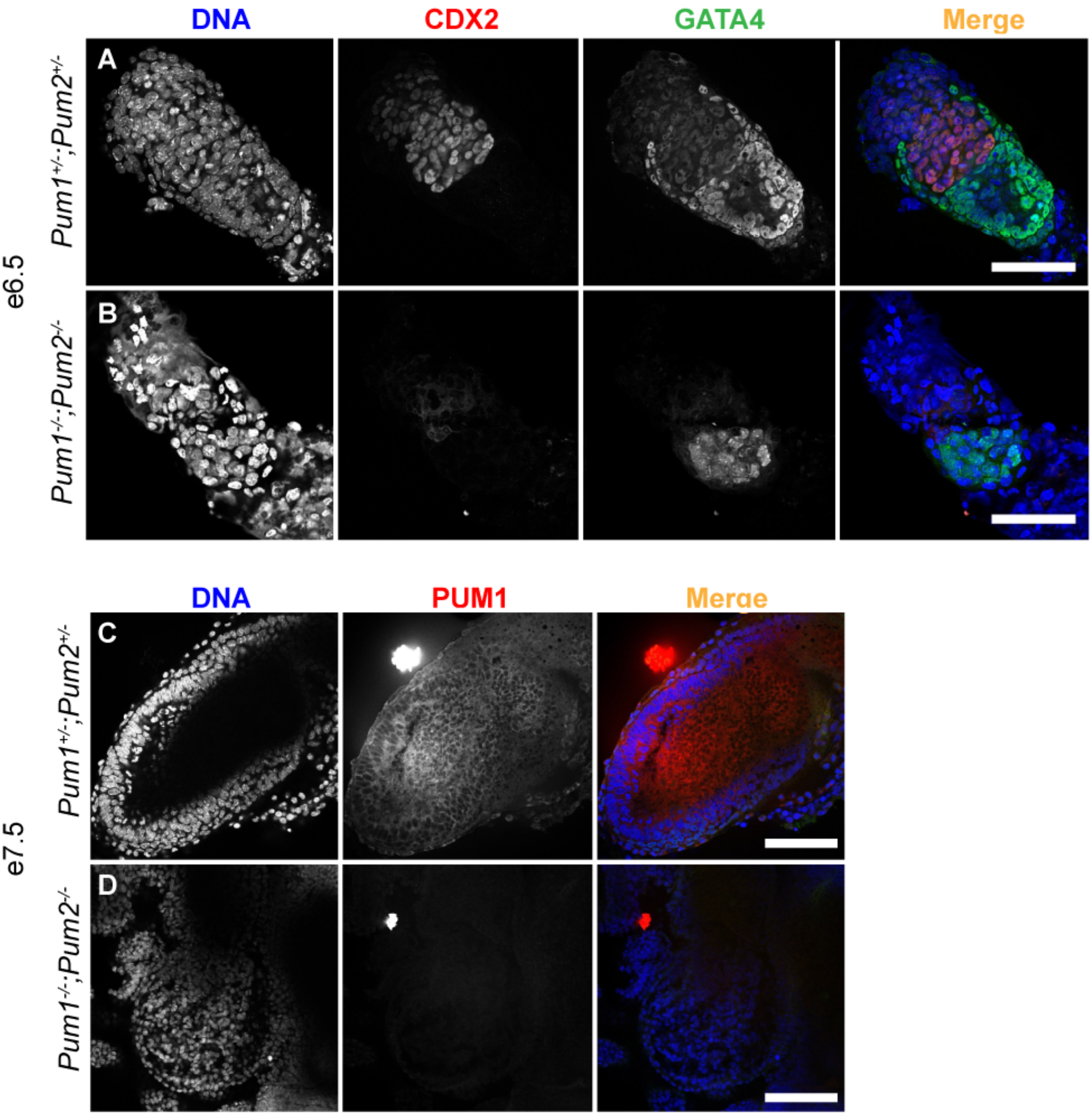

Supplemental Figure 7

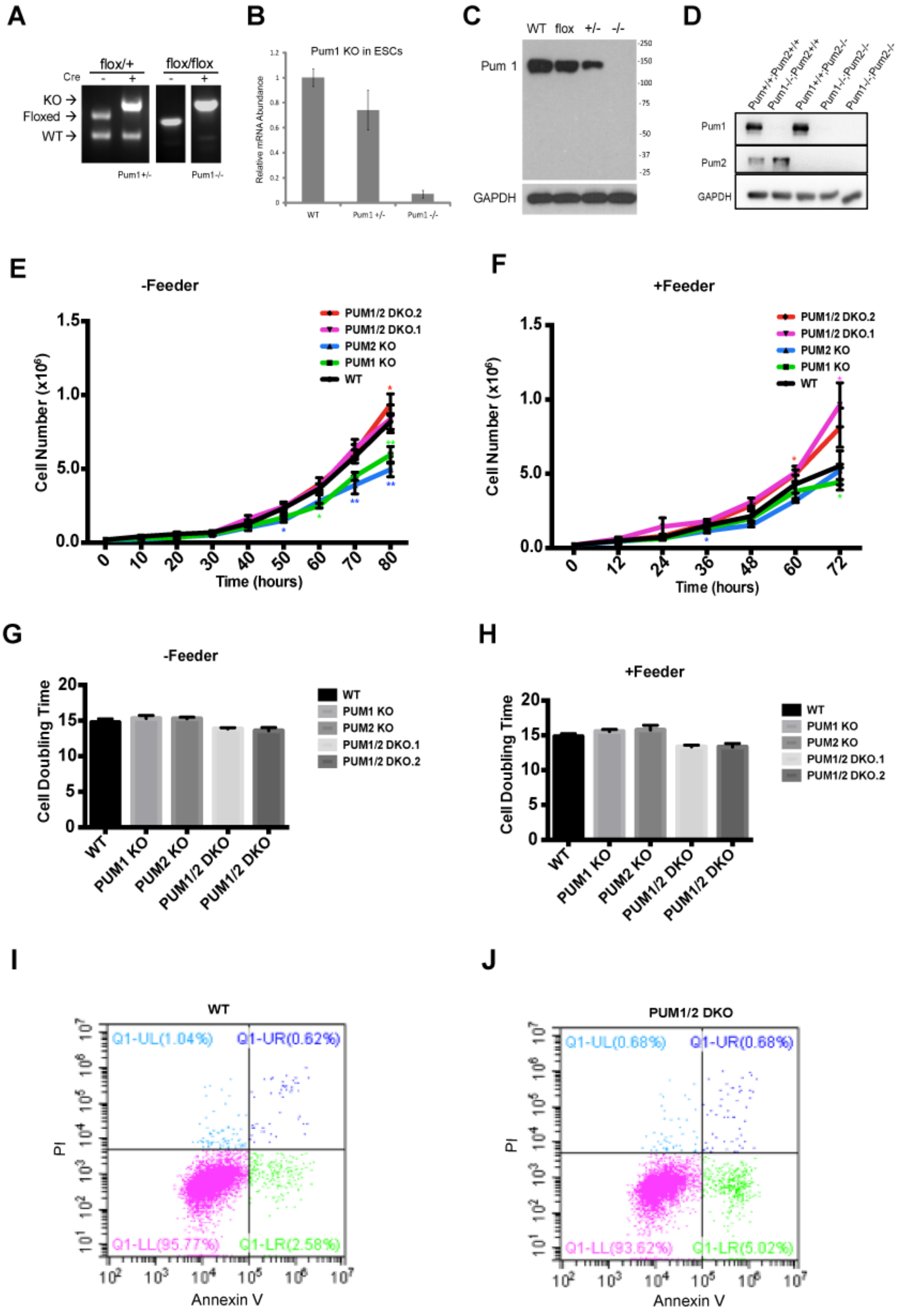

Supplemental Figure 8

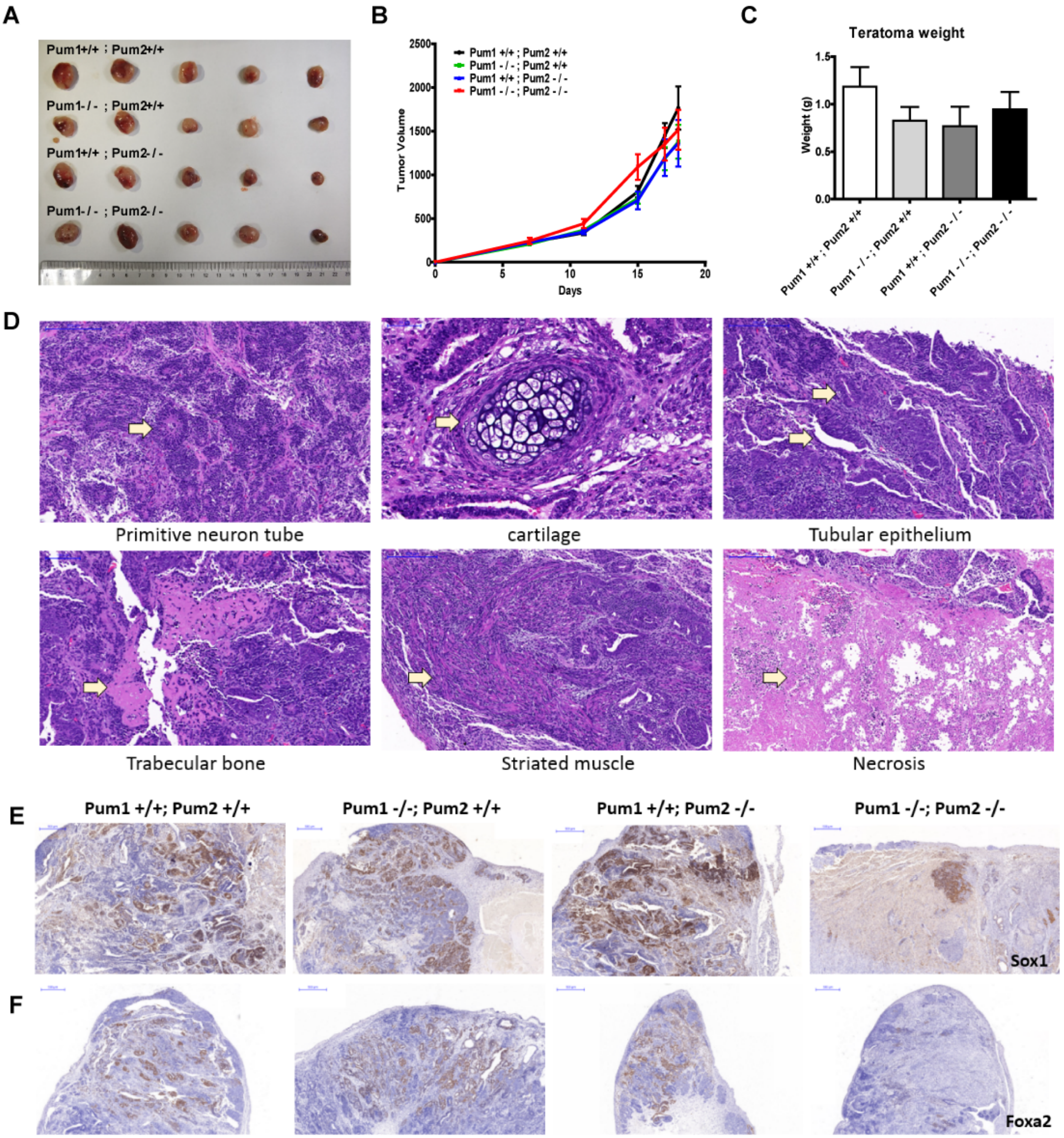

Supplemental Figure 9

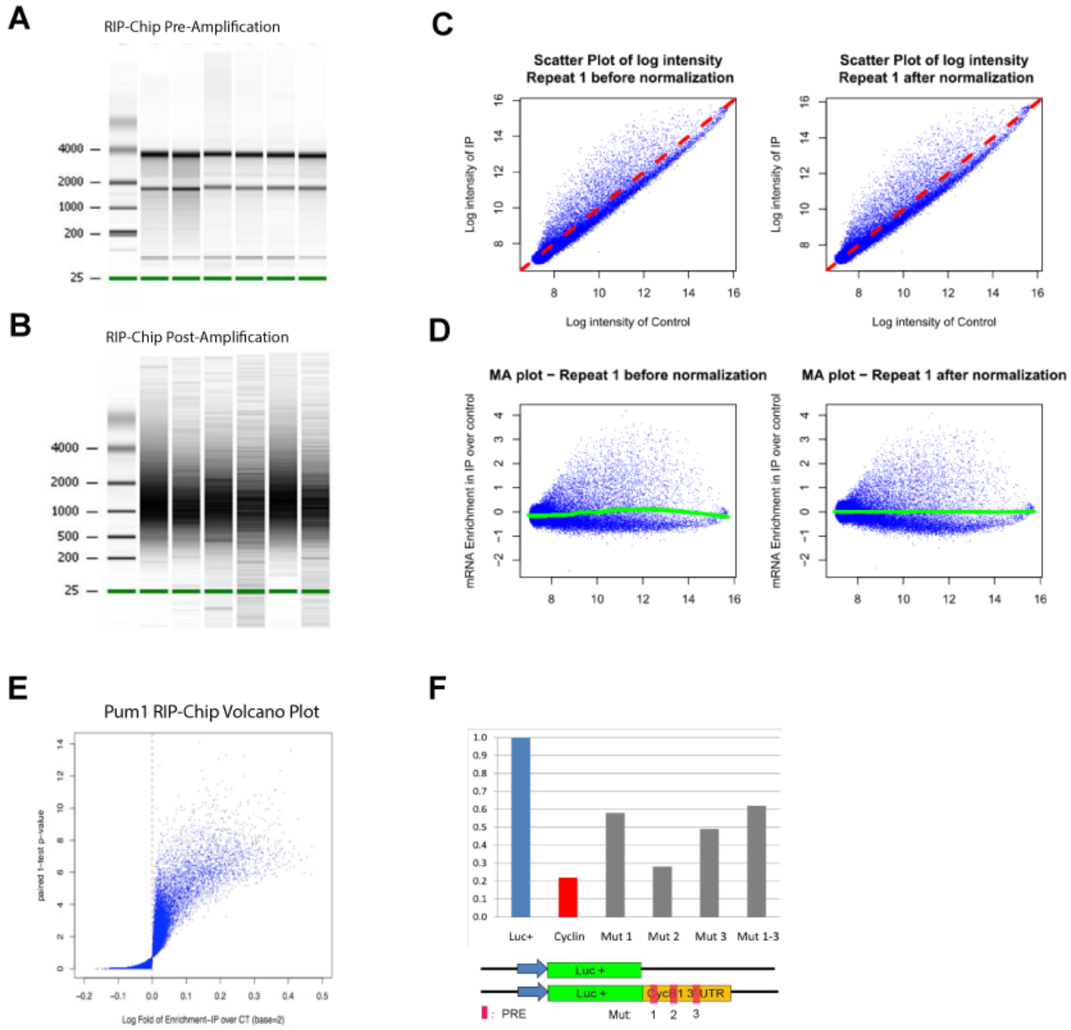

Supplemental Figure 10

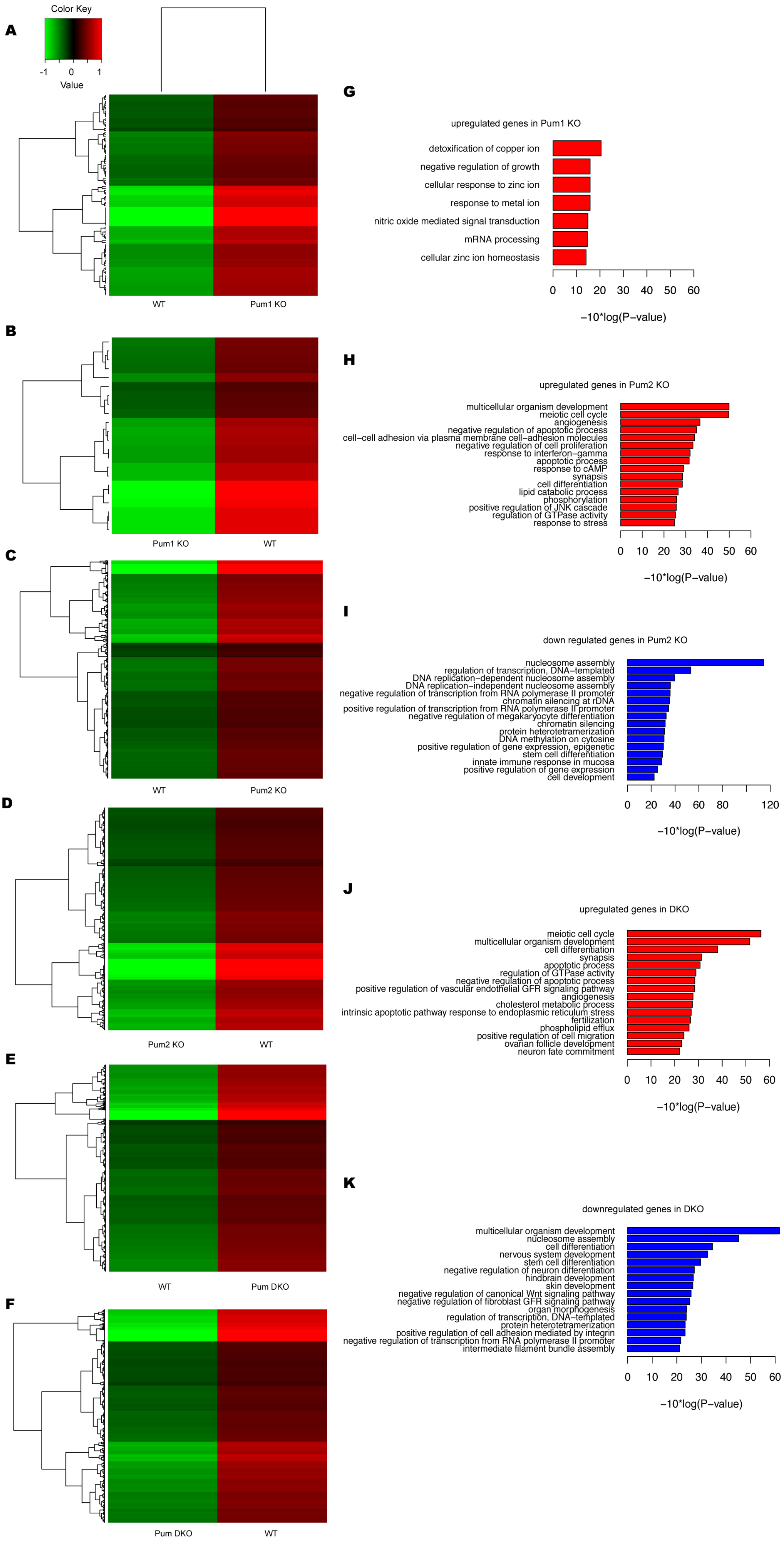

Supplemental Figure 11

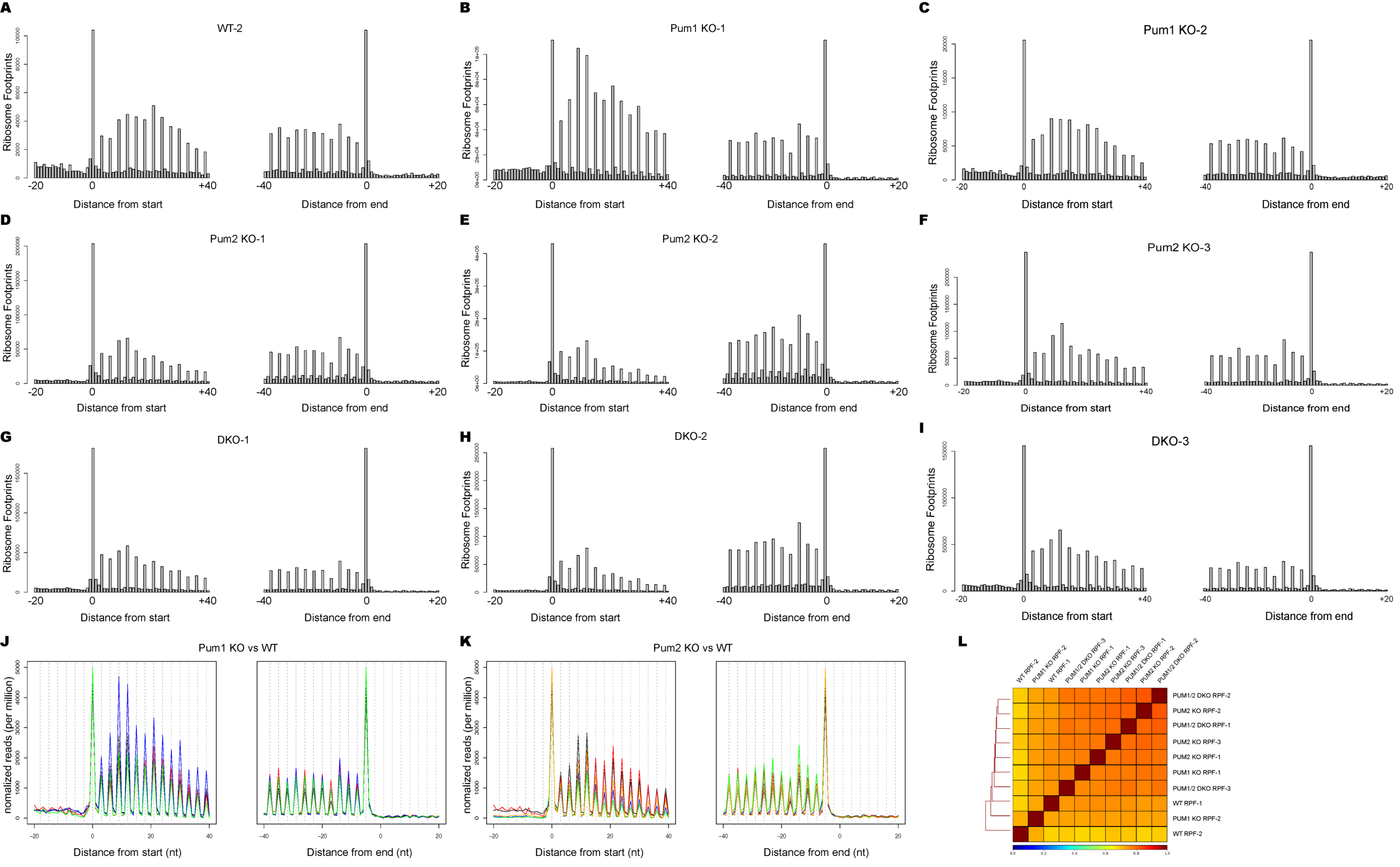
